# RNA promotes mitochondrial import of F1-ATP synthase subunit alpha (ATP5A1)

**DOI:** 10.1101/2024.08.19.608659

**Authors:** Aindrila Chatterjee, Michelle Noble, Thileepan Sekaran, Venkatraman Ravi, Dunja Ferring–Appel, Thomas Schwarzl, Heike Rampelt, Matthias W. Hentze

## Abstract

Most mitochondrial proteins are encoded by the nuclear genome, translated as precursor proteins in the cytosol and matured during directed import into the mitochondria ^1^. For many mitochondrial proteins this process is carefully regulated to meet demand and to avoid mitochondrial stress ^2,3,4^. Recently, mitochondrial F1-ATP synthase subunits have been found to interact with RNA across various eukaryotes. This includes genome wide RNA-interactome studies from yeast ^5–7^, fruit flies ^8,9^, plants ^10–12^, mice ^13–17^ and humans ^18–24^. To shed light on this unexpected observation, we determined the interacting cellular RNAs of ATP5A1 and the subcellular sites of interaction. Using RNA binding-deficient mutants of ATP5A1 and functional assays, we show that specific cytosolic RNAs bind ATP5A1 precursor proteins at the outer surface of mitochondria and promote their mitochondrial import both in vitro and in cellulo. These findings add an unexpected twist to understanding mitochondrial protein import and expand the growing list of riboregulated cellular processes.

## Main

Mitochondrial ATP synthase is the terminal complex of the oxidative phosphorylation (OXPHOS) chain, normally generating the bulk of cellular ATP. Highly sensitive to metabolic cues, its assembly as well as activity is altered in both physiological and pathological cellular transformations ^25–28^. Progressive deregulation of ATP synthase is a prevalent hallmark in the pathogenesis of cardiovascular, neurodegenerative, and metabolic disorders ^27^. Despite decades of extensive research into the structure, assembly and mechanism of action of the ATP synthase complex (Reviewed in ^29^), the cellular processes that alter its activity continue to be enigmatic.

Systems-wide RNA interactome capture experiments have identified F1-ATPase subunits ATP5A1 (α), ATP5B (β), and ATP5C1 (γ) as RNA binders in mammalian cells, but its biological significance remains unclear. To validate RNA binding in human cells (Huh7), we employed the complex capture (2C) assay that leverages RNA binding to silica-columns ^7^. We observed that following UV-crosslinking of RNA-binding proteins (RBPs) to RNA in living cells and 2C selection of lysates under highly denaturing conditions, ATP5A1, ATP5B, and ATP5C1, but not TOM20 (another mitochondrial membrane protein) or H3 (a DNA binder), co-purified with RNA (Fig. 1a, Extended Data Fig.1a). Notably, the association was lost when UV crosslinking was omitted or when the samples were digested with RNaseI prior to column purification (Extended Data Fig.1a). As an orthogonal approach, we immunoprecipitated ATP5A1 or ATP5C1, and tested their association with RNA by an end-labeling reaction with radioactive ATP and polynucleotide kinase (PNK). Both ATP5A1 (Fig. 1b) and ATP5C1 (Extended Data Fig. 1b) exhibited the characteristic smear indicative of RNA binding, and this smear was sensitive to RNaseI treatment. Collectively, these experiments validate ATP5A1, ATP5B, and ATP5C1 as bona fide RNA-binding proteins (RBPs) in human Huh7 cells. For deeper exploration, we focused on ATP5A1.

**Fig. 1:**
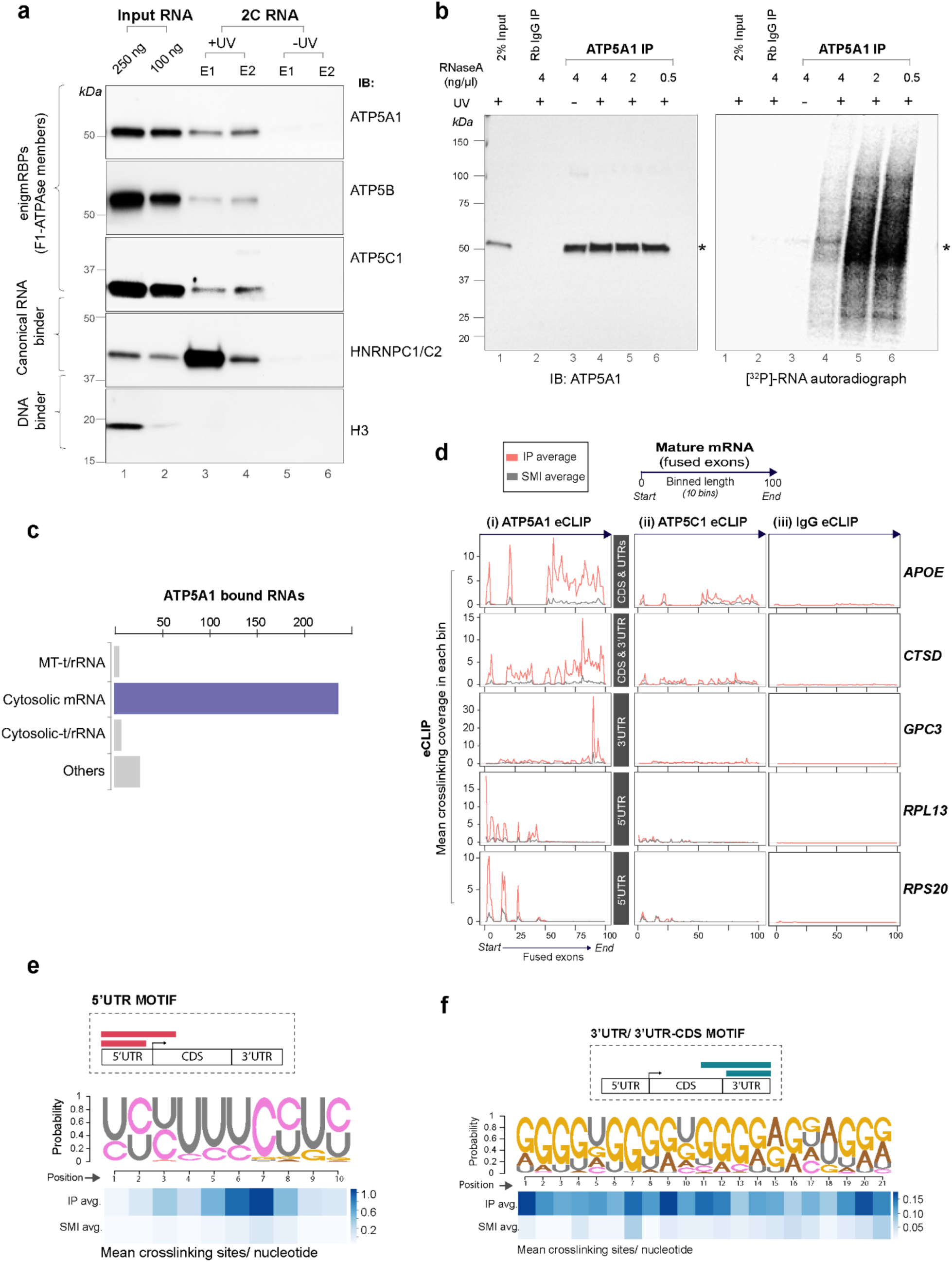
Mitochondrial F1-ATPase proteins bind RNA. a. Complex capture (2C) assay to detect proteins co-purified with RNA via immunoblotting. 50 µg of 2C RNA loaded, from UV-crosslinked or non-crosslinked controls. Lanes 1-2 represent pooled inputs from two independent experiments (E1, E2). HNRPC1/2 and H3 serve as positive and negative controls, respectively. b. Radioactive (^32^P) polynucleotide kinase (PNK) assay to visualise ATP5A1 association with RNA. Immunoblot (left) and [^32^P] RNA autoradiograph (right) after ATP5A1 IP are shown. Signal specificity confirmed by RNase A sensitivity. Asterisks = band of interest. c. RNA types significantly enriched in the human ATP5A1 eCLIP-seq dataset. DEWSeq analysis, exonic reads, mean IP count > 25, mean Log_2_ fold change ≥ 1, p.Adj ≤ 0.05, *n* = 3 biological replicates. d. Metagene plots, generated using an R/Bioconductor package called metagene2 (v1.16), show mean crosslinking signal coverage of eCLIP and size-matched input (SMI) controls for ATP5A1 on 6 representative mRNAs primarily bound at UTRs and/or CDS regions. mRNA lengths are binned (total 10) for comparing different RNA lengths, after exon fusion. Comparative eCLIP coverage plots for ATP5C1 (ii) and IgG (iii) are also displayed for the same mRNAs. *n =* 3 biological replicates e. Distinct sequence motifs recognised by ATP5A1 at 5’ and 3’ UTRs, **e-f**. Heat maps show probabilities of crosslinking site occupancy. *n =* 3 biological replicates. f. X

We next set out to identify the transcriptome-wide RNA targets of ATP5A1 using eCLIP-Seq^30^. We also included ATP5C1 in the analysis for comparative assessment and used IgG as a specificity control. The antibodies used for eCLIP-seq were rigorously validated using knockdowns, immunofluorescence and immunoprecipitation assays (data not shown). Application of the DEW-Seq analysis pipeline ^31^ for eCLIP data robustly identified significantly enriched regions of RNA binding across three replicate datasets (mean padj < 0.05 and mean IP crosslinking counts > 25) (Extended Data Table 1). In total, ATP5A1 binds 422 regions across 275 unique RNAs and binds to regions that are distinct from ATP5C1 or the IgG control (Fig. 1c, Fig. 1d). Surprisingly, only a small number of interacting RNAs are mitochondrial and ∼ 86% of the identified ATP5A1 targets are cytosolic mRNAs (236) (Extended Data Table 1, cluster analysis). Furthermore, the identified ATP5A1 targets share linear sequence motifs: a ∼10-nucleotide polypyrimidine (CU) motif located in 5’UTR regions (Fig. 1e) or a ∼20-nucleotide G-rich motif contained mostly in the 3’UTR/3’UTR-CDS region of bound RNAs (Fig. 1f). Interestingly, ATP5A1-bound RNAs display functional associations, with eukaryotic translation emerging as the largest cluster (Fig. 2a). The mRNAs within this cluster primarily correspond to those that harbor the conserved 5’ terminal oligopyrimidine (TOP) motif CU(C/U)UU(C/U)C ^32^. ATP5A1 exhibits robust binding to the TOP motifs within this subset of its targets, in line with the high similarity between the TOP motif (Fig. 2b) and the de novo identified motif for ATP5A1 (Fig. 1e). We confirmed the association of ATP5A1 with multiple targets by fRIP-qPCR (0.1% formaldehyde cross-linked RNA immunoprecipitation) from Huh7 cells, showing significant binding specificity with target mRNAs (GPC3, CTSD, RPS6, RPS20, RPL13) and exhibiting significantly higher enrichment than specificity controls (RPLP0 and ATP5C1) of similar RNA abundance (Fig. 2c). We also validated these targets using an imaging-based approach. We applied RNA-Protein proximity ligation assay (rnaPLA), using labeled antisense probes (biotin or DIG) against candidate RNAs and different antibodies against ATP5A1(mouse or rabbit). Again, we observed high specificity for the ATP5A1-RNA interactions in all tested combinations (Fig. 2d-e, Extended Data Fig. 1c), corroborating the previous fRIP-qPCR data.

**Fig. 2:**
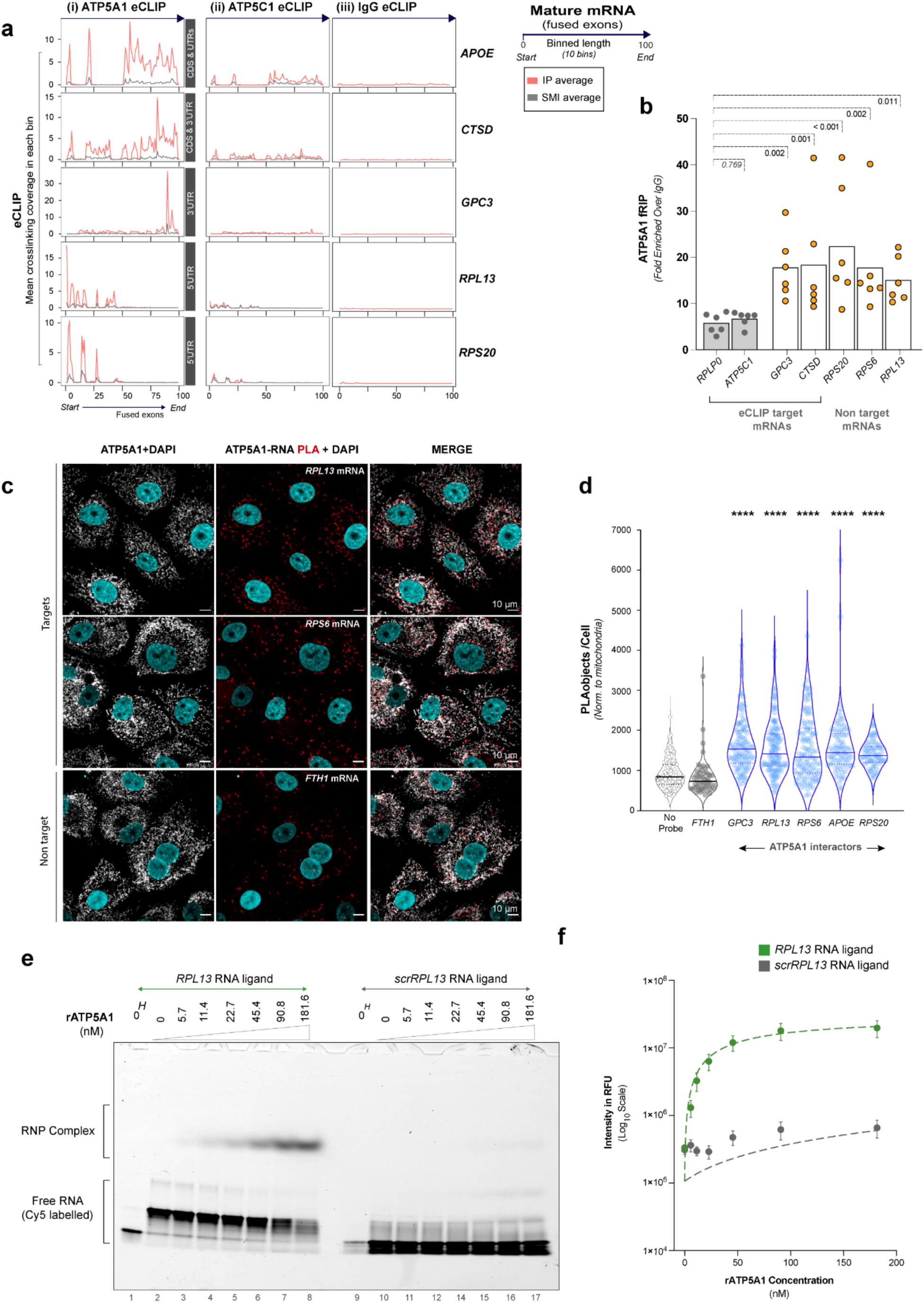
ATP5A1 binds specific functionally related mRNAs. a. Bubble plot shows functional enrichment analysis of ATP5A1-bound mRNAs in eCLIP. Colour shading, Log_10_ transformed Benjamini-Hochberg adjusted p-values (≤ 0.05). Bubble size, proportional to the number of genes represented in a pathway term, minimum = 5. b. ATP5A1 occupancy on TOP motif, present in majority of 5’UTR bound mRNAs. c. Formaldehyde crosslinked (0.1 %) RNA immunoprecipitation (fRIP) qPCR assay for endogenous ATP5A1. Enrichment for target (*GPC3, CTSD, RPS20, RPS6, RPL13*) or non-target (*ATP5C1, RPLP0*) mRNAs is calculated as ratio of percentage input recovery of ATP5A1 IP over IgG control IP. Ordinary two-way ANOVA performed against RPLP0, with Holm-Šidák’s post hoc test. *n* = 6 independent experiments. d. Representative confocal images of ATP5A1-RNA proximity ligation assay (rnaPLA) for target (*RPS6, RPL13*) and non-target (*FTH1*) mRNAs. ATP5A1 (white), rnaPLA signal (red) and nucleus (DAPI, cyan). e. ATP5A1 RNA interactions analysed by rnaPLA assay, for target mRNAs (*GPC3, RPL13, RPS6, APOE, RPS20),* non-target mRNA (*FTH1*) and no probe control. ATP5A1 was labelled with a rabbit polyclonal antibody. Cell size normalised by mitochondrial count, ∼ 50-100 cells per group, *n* = 4 independent experiments. Ordinary one-way ANOVA performed against no probe control, with Holm-Šidák’s post hoc test, ****p < 0.0001.

To gain further insight into the interactions of ATP5A1 with cytosolic mRNAs, we employed high resolution microscopy (Airyscan) to resolve ATP5A1-RNA interactions by RNA-PLA (Fig. 3a-b). We made three major observations. First, the majority of ATP5A1 signals localize to the mitochondria, representing the pool of mature protein. Second, the general pattern of mRNA-ATP5A1 interactions closely follows the mitochondrial network. And most importantly, these interactions occur at or in very close proximity to the outer mitochondrial membrane (OMM, stained by TOM70, Fig. 3a). This set of observations applies to all specific targets tested, which show robust enrichment over controls (Fig. 3b).

**Fig. 3:**
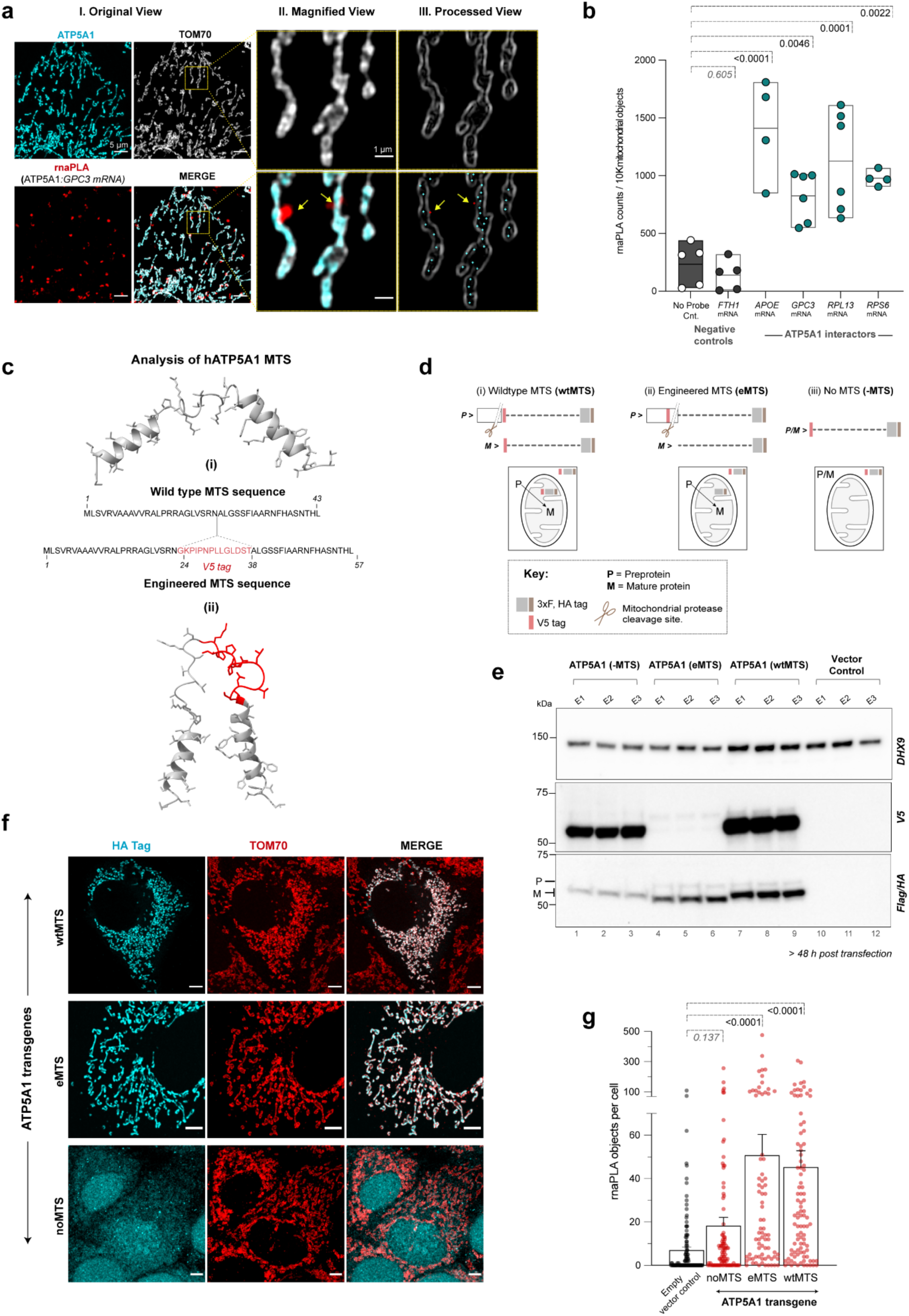
ATP5A1 pre-protein binds RNA at the outer mitochondrial membrane. a. Representative super resolution images (Airyscan SR, maximum intensity projection of z-stacks) from ATP5A1-*GPC3* rnaPLA assay. ATP5A1 (cyan), rnaPLA signal (red) and outer mitochondrial membrane (OMM) protein TOM70 (white) are shown. Processed view (III) shows OMM outline (white), alongwith centroids of signals from ATP5A1 staining (cyan) and rnaPLA signal (red). Yellow arrows indicate the site of ATP5A1 and *GPC3* mRNA interaction. Scale bars represent 5 µm (original view, I) or 1 µm (magnified view, II). b. Box plots of rnaPLA counts from Airyscan SR images for target (*APOE, GPC3, RPL13, RPS6*) and non-target (*FTH1)* mRNAs alongwith no probe control. Field of view differences have been normalised by identified mitochondrial objects using TOM70 staining. Each dot represents an image, collected from 3 independent experiments. Ordinary one-way ANOVA performed against no probe control, with Holm-Šidák’s post hoc test. c. AlphaFold2 predicted structure of mitochondrial targeting signal (MTS) with original sequence shown below, for (i) wild type hATP5A1 and (ii) its engineered version (eMTS) with integrated V5 sequence. d. Cartoon depicting protein behaviour of various MTS-containing ATP5A1 constructs. Mature (M) and pre-protein (P) versions of ATP5A1, with V5/Flag/HA tags are indicated for each construct. Different MTS versions, wt = original wild type, eMTS = engineered, no MTS = deleted construct. e. Representative confocal microscopy images of different MTS containing ATP5A1 constructs with C-terminal HA tag (cyan). Co-stained for mitochondria (TOM70, red) and nucleus (DAPI, blue). f. rnaPLA assay analysis for *RPS6* mRNA and V5 tag of different ATP5A1 transgenes. Non-mitochondrial (-MTS), pre-protein (eMTS), mature or pre-protein (wtMTS) containing ATP5A1 constructs. Data collected from average ∼ 110 cells per group. Ordinary one-way ANOVA performed against no probe control, with Holm-Šidák’s post hoc test. *n* = 3 independent experiments. g. Representative fluorescent electrophoretic mobility shift assay (EMSA) for bacterially expressed recombinant hATP5A1 (1-553, corresponding to ATP5A1 pre-protein) association with Cy-5 labelled RNA fragments-*RPL13,* target and its scrambled sc*RPL13,* control. *H* = RNA denatured at 95°C before gel loading. h. Non-linear fit of hATP5A1 association with target (*RPL13*) or non-target (sc*RPL13*) RNA ligands. Dots represent the mean ± SEM of 6 independent experiments, using 3 separate protein purifications, each used twice. Replicate values in Extended Data Fig. 2d.

Since ATP5A1 is translated in the cytoplasm as a pre-protein, we reasoned that the pre-protein of ATP5A1 might be the cytoplasmic mRNA interacting form. However, distinguishing between the precursor and mature ATP5A1 on a gel poses a challenge due to their minimal mass difference (∼2 kDa) ^33^. To address this issue, we generated an ATP5A1 variant with an engineered mitochondrial targeting signal (eMTS) that contains an internal V5 epitope tag within the disordered/flexible region of the original MTS (Fig. 3c). This tag facilitates selective identification of the pre-protein version, and improves the size differentiation between pre- and mature ATP5A1 in biochemical assays (Extended Data Fig. 2a-b). The eMTS (Fig. 3c, i) was carefully designed to avoid disrupting the overall MTS structure (Fig. 3c, ii) or function ^34,35^ (helical amphipathicity, Tom20 binding site and mitochondrial peptidase cleavage sites). As controls, we generated ATP5A1 variants with a wtMTS and without MTS (noMTS), both of which contain the V5 tag at the N-terminus of the mature protein (Fig. 3d). In addition, all versions of ATP5A1 were C-terminally tagged with 3xFlag-HA to identify all forms of ATP5A1 independent of their maturation status (Fig. 3d).

Confocal microscopy confirmed that eMTS-ATP5A1 shows similar mitochondrial localization as the wtMTS-ATP5A1, while the noMTS-ATP5A1 variant distributed throughout the cell (Fig. 3e). Consistent with the subcellular distribution, immunoblotting (Extended Data Fig. 2b-c) confirmed the MTS cleavage of both wtMTS- and eMTS-containing ATP5A1 transgenes, attesting to the appropriate maturation and mitochondrial targeting of the engineered versions of ATP5A1 except the noMTS control.

We then performed rna-PLA assays to score the interactions between transgenic versions of ATP5A1 and one of its specific RNA ligands, *RPS6* mRNA (Fig. 3f). While the rna-PLA with wtMTS-ATP5A1 replicated the enrichment seen with endogenous ATP5A1, we observed that loss of mitochondrial targeting (noMTS-ATP5A1) profoundly reduced the mRNA association. However, the pre-protein per se is fully competent for mRNA binding, as evidenced by the V5-tag-positive form of eMTS-ATP5A1 (Fig. 3f). As an orthogonal confirmation, we performed in vitro electromobility shift assays with recombinant ATP5A1 pre-protein and RPL13 ligand RNA or a scrambled version thereof (Fig. 3g-h, Extended Data Fig. 2d), demonstrating directly that the ATP5A1 pre-protein is capable of specific RNA binding. These data collectively show that the outer surface of mitochondria is a preferred site of interactions between ATP5A1 precursor proteins and ligand RNAs.

This scenario raised the intriguing question of whether RNA binding might influence ATP5A1 precursor import. To test this possibility, we conducted in vitro protein import assays with isolated human mitochondria that had been subjected to limited RNase A treatment or left as mock-treated controls (Fig. 5a). Labeled ATP5A1 precursor protein was generated by in vitro translation with 35S-methionine, and its mitochondrial import and processing was followed by gel electrophoresis over a 10 minute time course. Remarkably, ATP5A1 import demonstrated sensitivity to RNase treatment, as RNA removal led to a significantly reduced ATP5A1 import rate (Fig. 5b-c). Importantly, the import of the precursor of the mitochondrial matrix protein OTC (ornithine transcarbamylase) remained unaffected (Fig. 5b, 5d) under these conditions, demonstrating the specificity of the effect. Thus, RNA appears to facilitate the mitochondrial import of ATP5A1 *in vitro*.

To explore a possible role of RNA in mitochondrial import of ATP5A1 in cells, we next generated RNA binding-deficient mutants (RBdef) of ATP5A1. Towards this end, we leveraged mass spectrometric datasets ^20,21,36^ that indicate putative RNA-binding regions of RBPs (Fig. 4a). These datasets implicate a region of ∼100 amino acids of ATP5A1 located within the mature form of the protein and including the ADP-binding P-loop. We positioned mutations within this region aiming to affect RNA binding while preserving the overall structure and other functions of the protein. We selected three discrete mutants (Fig. 4a) from which we generated stable, inducible Huh7 cell lines that express C-terminally tagged versions of the ATP5A1 wildtype or MUT 1-3 (Fig. 4b). Comparative fRIP-qPCR analysis confirmed that MUT 1/2/3 exhibit significantly reduced binding to the specific RNA targets, while maintaining a similar degree of non-specific RNA interaction as the wild type (Fig. 4c).

**Fig. 4:**
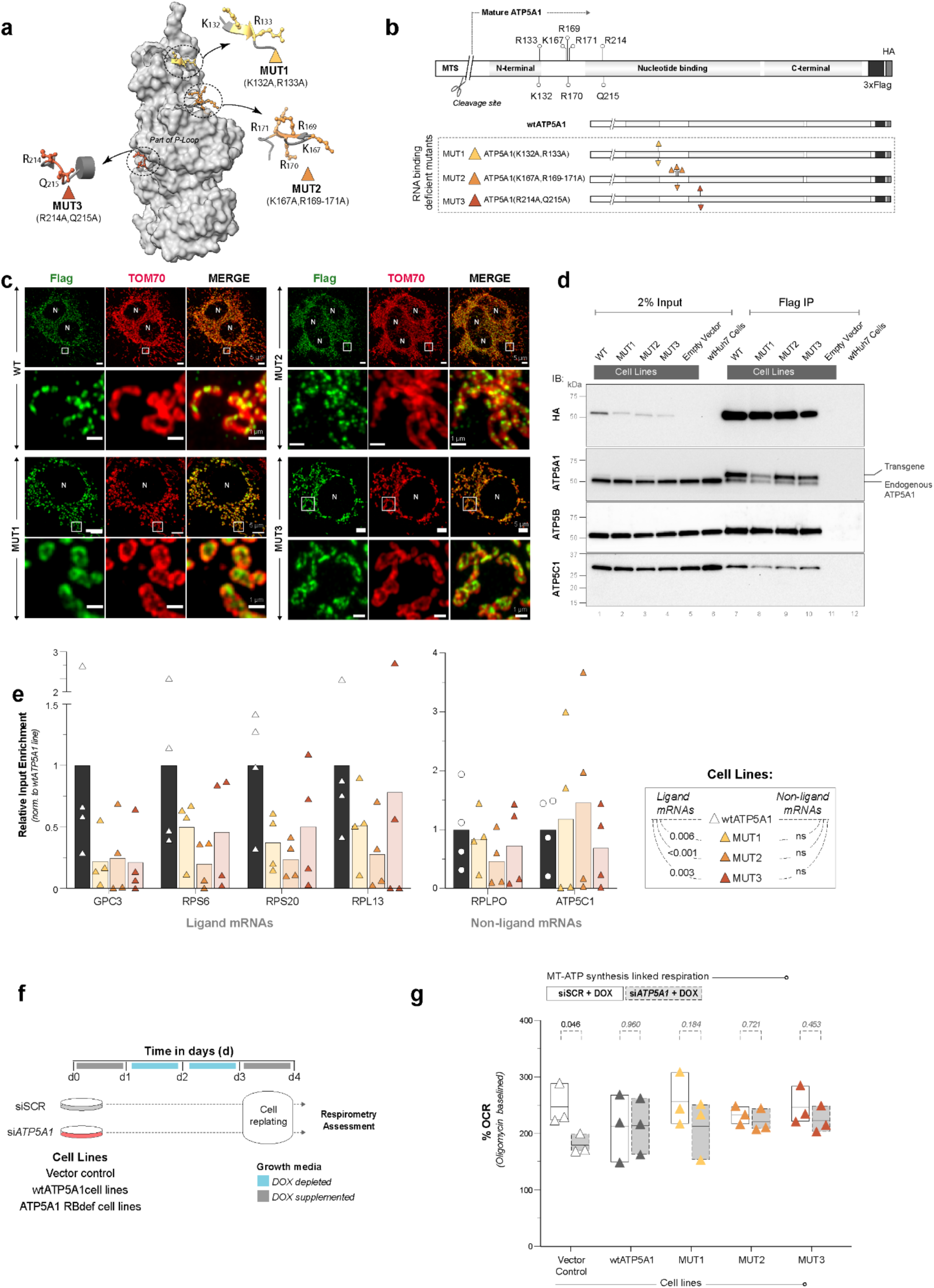
ATP5A1 has a broad RNA binding surface. a. Putative RNA binding residues mapped on ATP5A1 surface (ATP synthase alpha subunit, PDB: 1e79). MUT1-3 denote three RNA binding deficient (RBdef) mutants, designed on the basis of spatially distinct binding regions. b. RBdef mutation sites location with respect to ATP5A1 functional domains. C-terminal 3xFlag, HA tags were incorporated in transgenic constructs. c. fRIP-qPCR assay of C-terminally tagged (3xFlag,HA) ATP5A1 cell lines after Flag IP. Enrichment of target and non-target mRNAs in RBdef mutant cell lines (MUT1-3) were compared against wild type ATP5A1 line. Two-way ANOVA, Holm-Šidák’s post hoc test, significant P-values (<0.05) are indicated in figure. *n* = 4 independent experiments. d. Representative confocal images of different RBdef mutant cell lines of ATP5A1 (C-terminally 3xFlag,HA tagged) identified by Flag antibody (green) and co-stained for mitochondria (TOM70, red). N = nucleus. e. Immunoprecipitation (Flag) and western blotting of C-terminally tagged ATP5A1 wild type (wt) and RBdef mutant (MUT) cell lines. T = transgene, E = endogenous ATP5A1. f. Experimental design for assessment of mitochondrial ATP-linked respiration in vector control, wtATP5A1 and RBdef mutant cell lines (MUT1-3) after *ATP5A1*/SCR knockdown. g. % OCR (oxygen consumption rate) linked to mitochondrial ATP synthesis, deduced from Seahorse respirometry assays. Cell lines analysed after 96 hours of siSCR control (white boxes) or si*ATP5A1* knockdown (grey boxes) or, with DOX-induced transgene expression (scheme in panel f). *n* = 3 independent experiments.

Importantly, all three mutants localize to mitochondria (Fig. 4d) and bind their ATP synthase complex partners (ATP5A1, ATP5B, ATP5C1) (Fig. 4e), indicating proper protein folding and ATP synthase complex assembly. As a further test, we conducted functional Seahorse respirometry assays in transgenic cell lines where endogenous ATP5A1 was specifically depleted using siRNAs and the transgenic, siRNA-resistant variants (Fig. 4f) were induced for 24 hours. While ATP5A1 depletion causes the expected reduction in the oxygen consumption rate (OCR), both transgenic wtATP5A1 as well as the RNA binding mutants (MUT1-3) successfully restore the OCR. All of these experiments validate that MUT1-3 are not compromised in other key functions of ATP5A1 (Fig. 4g).

We then incorporated the eMTS design into MUT1-3 (Fig. 5e). When we transfected either wild type or MUT1-3 versions into Huh7 cells and followed their mitochondrial import over time, we found that all four versions were equally well imported into mitochondria 72 hours after transfection (Fig. 5f), in keeping with the full rescue of OCR by these proteins (Fig. 4g). However, at the earlier time points 48 hours and especially 24 hours after transfection, a clear distinction between the wild type and the MUT1-3 versions of ATP5A1 became apparent: the compromised RNA association of MUT1-3 significantly delays their mitochondrial import (Fig. 5f). To test the promotion of mitochondrial ATP5A1 import by RNA independently, we overexpressed distinct RNA ligands in Huh7 cells and monitored the effect on import of the wild type protein compared to the RNA binding-deficient MUT1(eMTS) 24 hours after transfection (Fig. 5h). Three key observations emerged: First, the RNA binding-deficient mutant was significantly less well imported, consistent with the findings in Fig 5f. Second, the import of wild-type, but not of MUT1, was enhanced by over-expression of specific ATP5A1 RNA ligands. Third, non-ligand RNAs or scrambled versions of the ligand failed to promote import, reflecting the specificity of the effect. Taken together, these results show that specific RNA ligands promote ATP5A1 import into the mitochondria of human Huh7 cells.

**Fig. 5:**
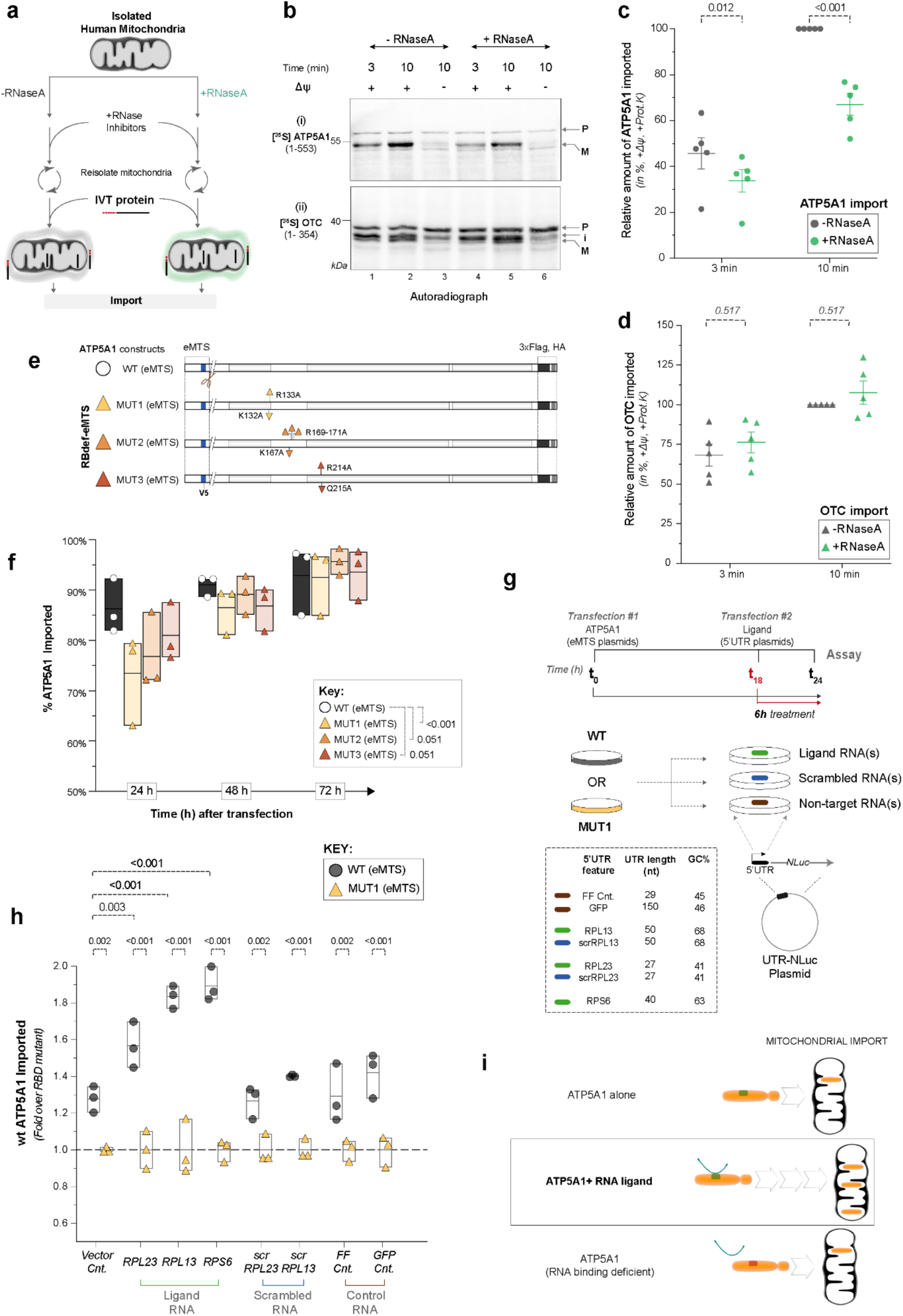
RNA promotes ATP5A1 import into mitochondria. a. Schematic representation of human mitochondrial import assays for in vitro translated (IVT) [^35^S] labelled protein(s). Freshly isolated human mitochondria were either subjected to limited RNaseA digestion or stored in the same buffer without treatment, as control. Digestion was stopped with RNasin before the import assay reaction and the mitochondria were re-isolated by centrifugation. b. Human mitochondrial import assay results are presented for ATP5A1 and OTC. **b,** Representative autoradiographs. P = Pre-protein, i = intermediate, M = mature/processed version in the matrix. OTC is dually processed (i and M) and both forms are considered in the quantification . **c-d,** quantifications from five independent experiments are displayed for the respective proteins. Samples were subjected to limited proteinase K treatment before loading. Δψ, mitochondrial membrane potential. c. X d. X e. Design of eMTS containing RBdef mutants (RBdef-eMTS). f. Time course of eMTS containing ATP5A1 constructs imported in Huh7 cells. RM one-way ANOVA for each time set, Holm-Šidák’s post hoc test, significant P-values (<0.05) indicated in figure. *n* = 3 independent experiments. g. Studying RNA ligand impact on ATP5A1 import: ATP5A1 expressed alone 18 hrs, NanoLuc plasmids with diverse 5’UTRs (different RNA ligands) expressed for the last 6 hrs pre-assay. **h**, Experimental scheme and 5’UTR sequences. **i**, Levels of wildtype ATP5A1 imported into mitochondria relative to its RNA binding deficient version, measured in the presence or absence of different RNA ligands. 2-way ANOVA, Holm-Šidák’s post hoc test, significant P-values (<0.05) indicated in figure. *n* = 3 independent experiments. h. X i. Schematic representation of riboregulation of ATP5A1 into mitochondria.

Recent work has elucidated that RNA binding by proteins can serve to riboregulate their respective functions or activities in cell biology, shedding light on the unexpected RNA-binding activity of a surprisingly large number of cellular proteins. This includes most glycolytic enzymes ^37,38^ such as GAPDH ^39–42^, Enolases ^37,38,43^, Phosphofructokinase ^37,44,45^ and Fructose-bisphosphate aldolase ^37,46^. Multiple TCA cycle enzymes ^47^ like, MDH2 ^48–51^, IDH1 ^52^, ACO1^53^, ACLY ^54^. Autophagy pathway proteins, LC3B ^55^, SQSTM1 (p62) ^21^. This report extends the biological scope of riboregulation to transmembrane transport of proteins. Although our data clearly show that direct and specific RNA binding to the ATP5A1 precursor protein enhances its mitochondrial import (Figs. 4 and 5), and that RNA binding preferentially occurs at or near the outer mitochondrial membrane (Fig. 3), we do not yet understand the mechanistic details that underlie RNA-facilitated mitochondrial import. Possibilities include RNA-mediated enrichment of the import substrate near the mitochondria, an RNA chaperoning function that promotes engagement of the import substrate with the import machinery, or direct RNA bridging of import substrate and import machinery. It will also be interesting to determine whether this process is regulated, for example in response to increased or decreased needs for mitochondrial ATP synthesis. Since dozens of transmembrane proteins, extracellular proteins and organelle-resident proteins have been found to bind RNA, our findings addressing the RNA-binding activity of ATP5A1 may foreshadow a much broader role of riboregulation in transmembrane transport.

## Acknowledgements

We thank S. Sahadevan (EMBL, Heidelberg) for his expert input on bioinformatic analyses and I. Perschil (University of Freiburg) for technical support . We acknowledge the invaluable expertise and support of EMBL Heidelberg core facilities for advanced light microscopy (M. Gunkel and A. Halavatyi), genomics, protein expression and purification (A. Boergel). A.C. is thankful to C. Tischer (Center for Bioimage Analysis, EMBL) for training on imaging data analysis. We are grateful to members of the Hentze laboratory for their critical discussions and feedback. H.R. is supported by Deutsche Forschungsgemeinschaft (DFG, German Research Foundation) for 2848 – project ID 401510699 and Germany’s Excellence Strategy CIBSS – EXC-2189 – project ID 390939984. M.W.H. acknowledges funding from the Manfred-Lautenschläger Stiftung and the DFG CRC1550 Molecular Circuits of Heart Disease.

## Extended Data Figures and Figure legends

**Extended Data Fig. 1:**
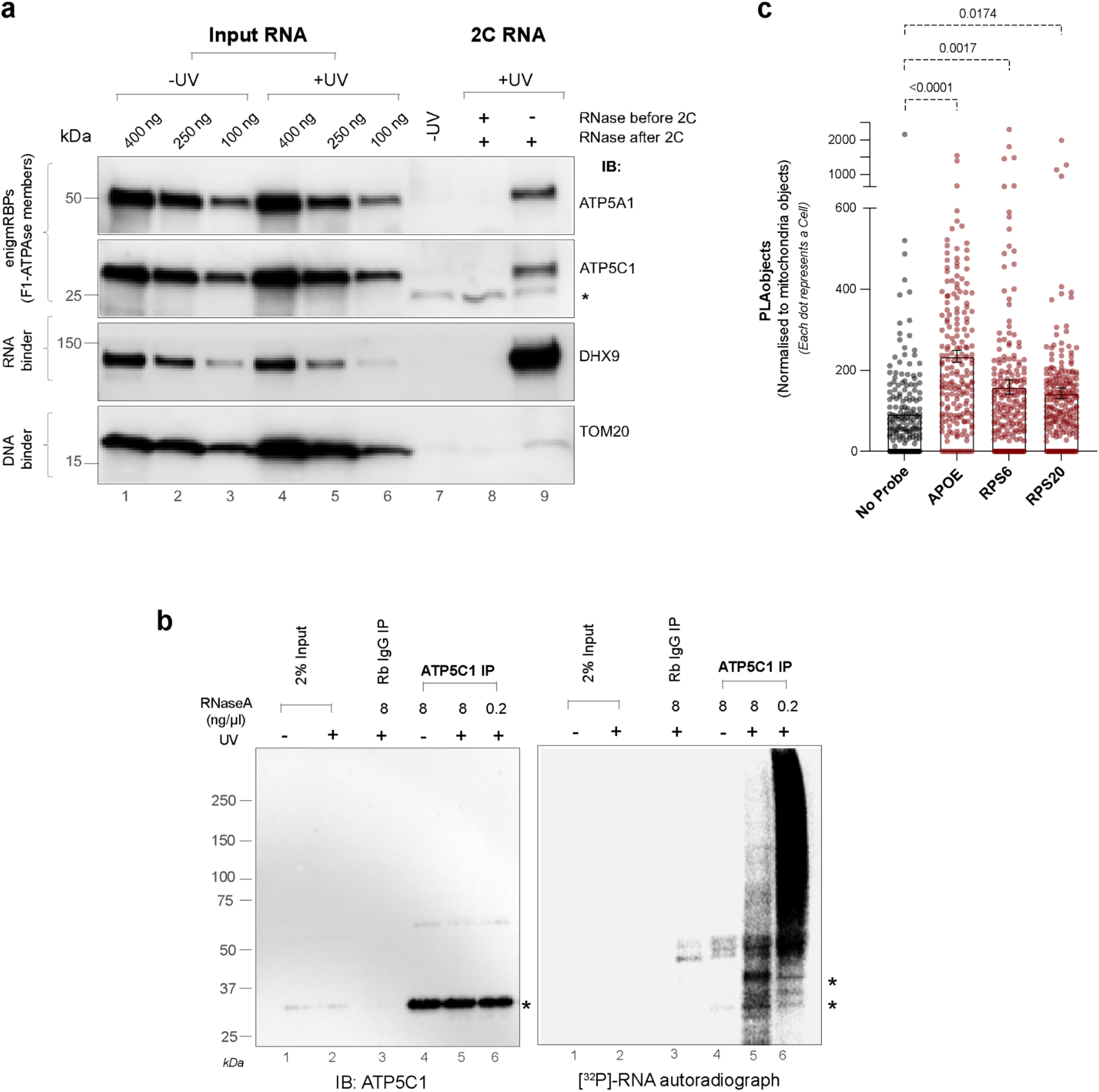
Mitochondrial F1-ATPase proteins bind RNA. a. Complex capture (2C) assay to detect proteins co-purified with RNA via immunoblotting. 45 µg of 2C RNA was loaded from UV crosslinked or non-crosslinked controls. DHX9 and TOM20 serve as positive and negative controls, respectively. 500 U RNase I was used whenever mentioned. Asterisks = band of interest. b. Radioactive (^32^P) polynucleotide kinase (PNK) assay to visualise ATP5C1 association with RNA. Immunoblot (left) and [^32^P] RNA autoradiograph (right) after ATP5C1 IP are shown. Signal specificity confirmed by RNase A sensitivity. Asterisks = band of interest. c. ATP5A1 RNA interactions analysed by rnaPLA assay, with antisense (DIG-labelled) probe against target mRNAs (*APOE, RPS6, RPS20)* and no probe control. ATP5A1 was labelled with a mouse monoclonal antibody. Cell size normalised by mitochondrial count, ∼ 192 - 214 cells per group, *n* = 4 independent experiments. Ordinary one-way ANOVA performed against no probe control, with Holm-Šidák’s post hoc test, p-values are mentioned in figure.

**Extended Data Fig. 2:**
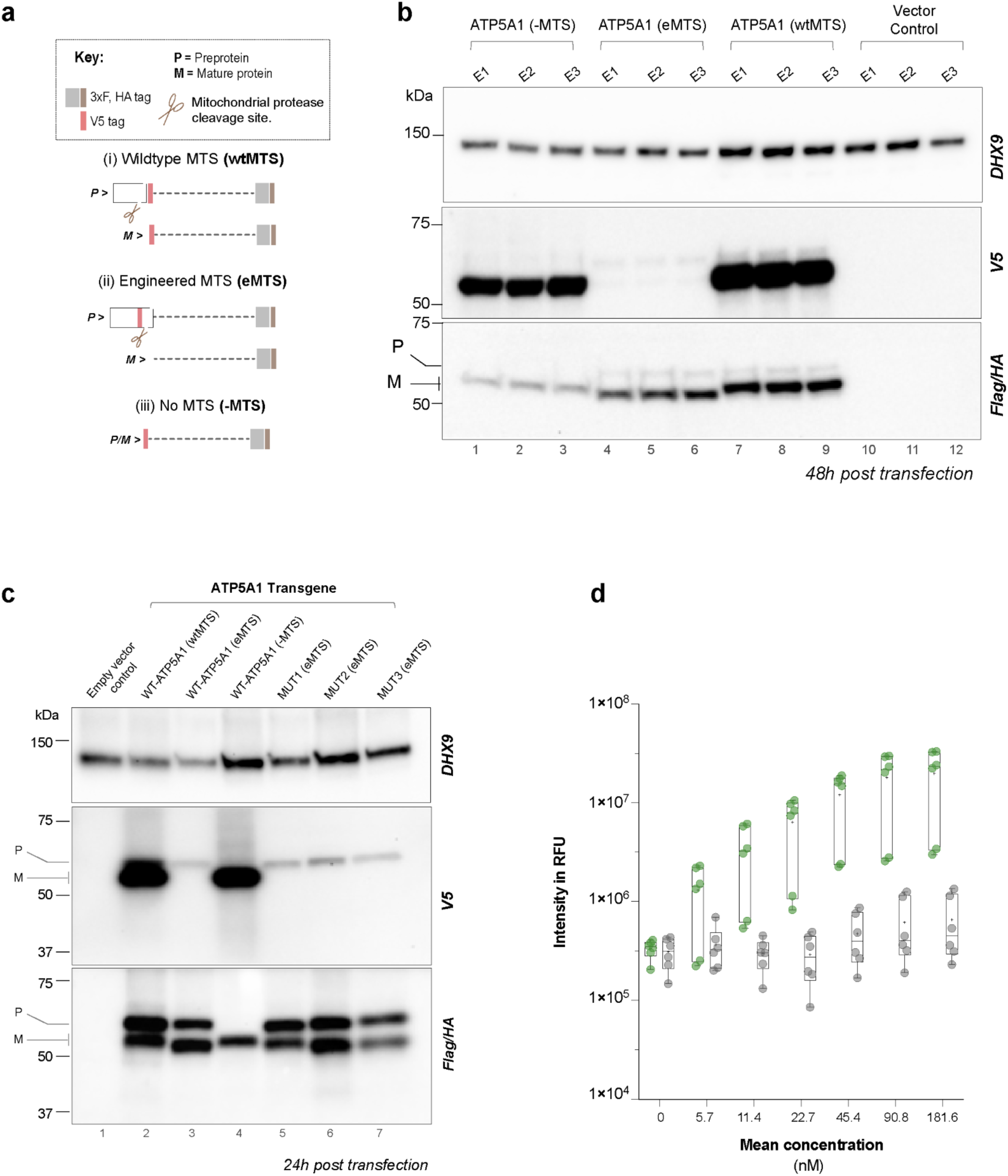
Mitochondrial F1-ATPase proteins bind RNA. a. MTS-containing ATP5A1 transgenic constructs. Mature (M) and pre-protein (P) versions of ATP5A1, with V5/Flag/HA tags are indicated for each construct. Different MTS versions, wt = original wild type, eMTS = engineered, no MTS = deleted construct. b. Western blot assessment mitochondrial targeting signal (MTS) cleavage patterns of various transgenic ATP5A1 constructs, **b-c.** Non-mitochondrial (-MTS), pre-protein (eMTS), mature or pre-protein (wtMTS) containing ATP5A1 constructs. Mature (M) and pre-protein (P) versions of ATP5A1 are indicated in the blot. **c,** eMTS containing ATP5A1 RBdef mutant (MUT1-3) transgenes. c. X d. In vitro association of hATP5A1(bacterially expressed) with target (*RPL13, green*) and non-target (sc*RPL13, grey*) RNA ligands.

## Methods

### Statistics and reproducibility

Experimental replicates are mentioned in the figures legends. GraphPad Prism 10 (Version 10.0.2) software was used for statistical data analysis, unless otherwise mentioned. Bar graphs display mean and all data points. Representative western blots are shown for experiments which have been repeated minimum three times. Experimenters did not blind, randomise or statistically pre-determine experimental sample size.

### Cell Culture and Maintenance

Huh7 (human, male) cells were cultured in low glucose DMEM base media (Sigma #D5523-10L)-with 5 mM Glucose, 1.25 mM Pyruvate, 2 mM L-Glutamine (Sigma #G7513), 10 % heat inactivated FBS (Thermo #10270106), 100 U/ml PenStrep (Thermo # 15140122). Unless otherwise mentioned, all reagents were purchased from Sigma. Cells were grown at 37°C with 5 % CO_2_, split at 90-95 % confluency using TrypLE express (Gibco #12604-013).

Huh7 Flp-In T-Rex host cells, as described by {Horos et al. in 2019}, were maintained with an additional 5 µg/ml of blasticidin (Invivogen #ant-bl-05) and 100 µg/ml of zeocin (Invivogen #ant-zn-05). Stable cell lines generated in this background were cultured with 5 µg/ml of blasticidin and 200 µg/ml of hygromycin B (Invivogen #ant-hg-1). To minimise background expression of the transgene, cell lines were cultured in the presence of 10% tetracycline-free FBS (PAN Biotech #P30-3602). Transgene expression was induced with 100 ng/ml doxycycline (Sigma #D3447-500MG) for a minimum 16 hrs.

HEK293T cells were cultured in DMEM media with 4.5 g/L Glucose (Sigma #D6429, or Gibco #21969 supplemented with 2 mM L-glutamine (Sigma #G7513)), supplemented with 10% FBS (Anprotec #AC-SM-0142) and 100 U/ml PenStrep (Gibco #15140).

### Buffers

#### Biochemistry

Unless otherwise mentioned, chemicals were purchased from Sigma. They were sterile filtered with a 0.22 µm filter before use. Wherever mentioned, DTT was always freshly added to the buffers.

- HMGN buffer: 20 mM HEPES pH 7.5, 150 mM NaCl, 2 mM MgCl_2_, 0.5% NP-40 and 10% glycerol.
- PBST-0.05% buffer: 0.05 % Tween20 in 1x PBS.
- RIPA buffer: 50 mM Tris-Cl pH7.5, 150 mM NaCl, 1% NP-40, 0.5% sodium deoxycholate and 0.1% and SDS.
- RIPA-HS buffer: 50 mM Tris-Cl pH7.5, 500 mM NaCl, 1% NP-40, 0.5% sodium deoxycholate, 0.1% SDS.
- PNK buffer: 50 mM Tris-Cl pH7.6, 50 mM NaCl, 0.5% NP-40, 10 mM MgCl_2_.
- Nuclease digestion buffer (NDB): 50 mM Tris-Cl pH7.5, 50 mM NaCl, 2.5 mM MgCl_2_, 1 mM CaCl_2_, 1 mM DTT.
- FastAP Buffer: 10 mM Tris-Cl pH7.5, 100 mM KCl, 5 mM MgCl_2_, 0.02 % Triton X-100.
- No salt wash Buffer (NSWB): 20 mM Tris-Cl pH7.5, 10 mM MgCl_2_, 0.2 % Tween20.
- T4 Ligase buffer: Purchased as 10x stock from NEB (#B0202S), used at 1x concentration.
- Streptavidin running buffer: 50 mM Tris-HCl pH8.0, 500 mM NaCl, 10% Glycerol.
- Streptavidin elution buffer: 50 mM Tris-HCl pH8.0, 500 mM NaCl, 10% Glycerol, 20 mM Desthiobiotin.
- SEC buffer: 10 mM Tris-HCl pH8.0, 150 mM NaCl, 10% Glycerol.

#### Immunohistochemistry

- Fixation buffer: 4 % Formaldehyde (16% methanol free stock, Thermo #28906) diluted in PBS.
- Permeabilization buffer: 0.2 % Triton X-100 in 1x PBS.
- PBST-0.02% buffer: 0.02 % Tween20 in 1x PBS.
- Immunofluorescence (IF) buffer: 2.5 % BSA, 0.02 % Tween20 in PBS.

#### RNA-Protein PLA

- Probe Hybridisation Buffer: 50 % deionized formamide, 5x SSC, 1x Denhardt’s reagent, 0.1 % (v/v) Tween 20, 0.1 % (v/v) CHAPS, 5 mM EDTA, 1 mg/ml of RNase free tRNA, (baker’s yeast), 100 ug/ml Heparin. All reagents were procured from Sigma.
- Duolink wash buffers A & B: The buffers are part of the Duolink in situ wash buffers fluorescence set (Sigma #DUO82049). They were reconstituted as instructed and stored at 4°C.

### UV crosslinking and cell lysate preparation

Huh7 cells at 90-95% confluency, were placed on ice, media removed and rinsed twice with pre chilled PBS. They were crosslinked in culture plates, on ice at UV 254 nm wavelength and 150 mJ/cm^2^ (∼ 40s) energy (Stratalinker 1800 UV crosslinker). Immediately afterwards, cells were lysed in either HMGN or RIPA buffer, supplemented with supplemented with RNase Inhibitor (Ambion) and EDTA-free protease inhibitors (MERCK #04693132001 Roche). Chromatin was sheared by sonication using either a Branson tip sonicator (*Settings:* 3 cycles, 10s ON, 10s OFF, 50% duty and Output 4) on ice or a Bioruptor water bath (*Settings:* High; 10 cycles, 10s ON/ OFF) at 4°C. Samples were cleared by centrifugation at 10,000 g for 10 mins at 4°C. Protein amount was estimated using Pierce 660 nm protein assay reagent (Thermo #22660). Snap frozen aliquots were stored at -80°C for up to 2 months and used in various assays.

### Knockdowns

Knockdowns were performed in Huh7 cells at 40 % confluency. We used Lipofectamine RNAiMAX Transfection Reagent (Invitrogen #13778075) and followed the manufacturer’s protocols. Pre-validated Silencer Select siRNAs (LNA modified) were purchased from Thermo. Specific siRNAs used were: si*ATP5A1* (Thermo # s1769 and s1768), si*ATP5C1* (Thermo # s189 and s191), si*ATP5B* (Thermo # s1773 and s1775). siRNAs and the RNAiMax reagent were prediluted in Opti-MEM reduced serum medium (Gibco #31985062) separately, before mixing. After confirming knockdown efficiencies of individual siRNAs (si*ATP5A1*, si*ATP5C1*, si*ATP5B*) at 10 nM concentration, we pooled the two siRNAs (10 nM each) for all knockdown experiments. For controls, we used pooled siSCR Controls: Silencer Select Negative Control No.1 (Thermo #4390843) and Silencer Select Negative Control No.2 (Thermo #4390846) at the same concentrations. Cells were cultured in antibiotics free low glucose DMEM, initially without serum (for the first 5-6 hrs), which was later supplemented to 10 % final concentration in fresh media. Based on trial experiments, we obtained the highest knockdown efficiency after 96 h of siRNA treatment.

### Transfections

Transfections were performed in Huh7 cells at 60 % confluency, using Lipofectamine 3000 (Thermo #L3000015) reagent, following recommended protocols. Cells were cultured in antibiotics free low glucose DMEM for minimum 24 h before assay.

### ATP5A1 pre-protein expression in *E.coli*

Full length hATP5A1 pre-protein was cloned in the petM22 vector with a c-terminal Strep-tag II tag and expressed in *E.coli* BL21 Rosetta (DE3). Bacterial induction followed the same steps as mentioned in Noble et al.^1^

#### hATP5A1 pre-protein sequence

MLSVRVAAAVVRALPRRAGLVSRNALGSSFIAARNFHASNTHLQKTGTAEMSSILEERILGADTSVDLEETGRVLSIGDGIARVHGLRNVQAEEMVEFSSGLKGMSLNLEPDNVGVVVFGNDKLIKEGDIVKRTGAIVDVPVGEELLGRVVDALGNAIDGKGPIGSKTRRRVGLKAPGIIPRISVREPMQTGIKAVDSLVPIGRGQRELIIGDRQTGKTSIAIDTIINQKRFNDGSDEKKKLYCIYVAIGQKRSTVAQLVKRLTDADAMKYTIVVSATASDAAPLQYLAPYSGCSMGEYFRDNGKHALIIYDDLSKQAVAYRQMSLLLRRPPGREAYPGDVFYLHSRLLERAAKMNDAFGGGSLTALPVIETQAGDVSAYIPTNVISITDGQIFLETELFYKGIRPAINVGLSVSRVGSAAQTRAMKQVAGTMKLELAQYREVAAFAQFGSDLDAATQQLLSRGVRLTELLKQGQYSPMAIEEQVAVIYAGVRGYLDKLEPSKITKFENAFLSHVVSQHQALLGTIRADGKISEQSDAKLKEIVTNFLAGFEAGSG*

#### Protein purification

Bacterial pellet was resuspended in 100 mL of Strep running buffer, supplemented with 5 mM MgCl2, 10 µL of SM nuclease (0.25 mg/mL), and EDTA-free protease inhibitors (MERCK #04693132001 Roche) were added. Cells were lysed with a microfluidizer (five rounds) followed by five cycles of 30-second sonication (50% output cycle and stage 5 intensity). Lysates were clarified by centrifugation at 35,000 rpm for 30 minutes at 4°C using a Ti45 rotor in a Beckman UC centrifuge. Supernatant was loaded onto a 5 mL Strep-tactin superflow column (IBA), pre-equilibrated with Strep running buffer, at a flow rate of 1 mL/min. The column was washed with a Strep running buffer until a stable baseline was achieved. ATP5A1 protein was eluted using a step gradient to 100% Strep elution buffer. The protein was loaded onto an SEC superdex s200 increase 10/300 column at 0.6 ml/min flow rate, equilibrated with SEC buffer.

### Seahorse Respirometry profiling of cells

Cellular respiration was assessed using the Seahorse XFe96 Extracellular Flux Analyzer (Agilent Technologies), following manufacturer and published protocol ^2^. In brief, 50,000 Huh7 cells were seeded into assay microplates the night before. The next day, cells were incubated in 160 µl of Seahorse XF DMEM Medium, pH 7.4 (Agilent #103575-100) at 37°C in a CO2-free incubator for 1 hour before the assay. The XFe96 analyzer measured the oxygen consumption rate (OCR) over time after sequential addition of respiratory chain inhibitors-oligomycin, FCCP, rotenone and antimycin A. All inhibitors were procured from Sigma, dissolved in DMSO and used in final assay concentrations of 1 µM. Post-assay, media was removed, cells were washed with PBS, and lysed for protein estimation. Data were normalised using the Pierce660 (Thermo #22662) total protein assay.

### Cloning

The wtMOF-sfGFP; MTS-wtMOF-sfGFP; MTS-MOF(ΔNLS1)-sfGFP, MTS-MOF(ΔNLS1,E350Q)-sfGFP; sfGFP and MTS-sfGFP sequences were cloned into pcDNA5/FRT/TO/NN vector backbone. This backbone was derived from pcDNA5/FRT/TO vector (Life Technologies) with modified MCS. All other features of the original vector were retained. These plasmids were used to generate Huh7 cell lines using the Flp-In T-Rex technology (Life Technologies). They were co-transfected with pOG44 (expresses Flp recombinase) into host Huh7 Flp-In T-Rex cells ^3^ to generate stable inducible cell lines. We synthesized ATP5A1, ATP5B, and ATP5C1 cDNA sequences using GeneArt Strings DNA Fragments (Thermo). Some long GC-rich stretches were modified (synonymous) for synthesis without altering protein sequences. Synonymous mutations were also introduced to make the transgenes insensitive to siRNAs used in this paper.

### Immunoblotting

Cell lysis, chromatin shearing, sample clarification and protein estimation adhered to the previously outlined protocol (without crosslinking step). For regular western blots, 10 µg RIPA lysates, and for IP/co-IP, 30-40% eluates were used. Samples were mixed with 1x LDS sample buffer (Invitrogen #NP0007), reduced with 1x TCEP (Sigma #646547), and heated at 70°C for 10 mins before gel loading. Typically, 4-15% Criterion TGX Precast Midi Protein gels (Bio-Rad) were run in 1x Laemmeli buffer and transferred via Trans-Blot Turbo Transfer System (Bio-Rad) to PVDF/nitrocellulose membranes. Membranes were blocked and probed with antibodies in 5% non-fat milk in PBST-0.05%. Standard protocols were followed for blocking (1 hr, room temperature), primary antibody (overnight, 4°C), and HRP-conjugated secondary antibody incubations (1 hr, room temperature). After antibody incubations, membranes were washed 3x with PBST-0.05% (5 mins per wash, room temperature). Chemiluminescence signals were developed using Pierce ECL reagents (Standard, #32109 or high sensitivity Femto, #34096) and imaged in the ChemiDoc imaging system (Bio-Rad).

### Complex capture assays (2C & 2C^2^)

#### 2C Assay

Standard complex capture (2C) assay was performed with modifications to the method described before ^4^. Briefly, Huh7 cells were crosslinked and lysed in RIPA lysis buffer, as described above. 1 mg frozen (maximum 1 month old) lysates were used for 2C RNA preparation in each Zymo-Spin IIICG column (Quick-RNA MiniPrep kit, Zymo #R1055). The published protocol was followed for the rest of the assay. 40-60 µg 2C-RNA was used for immunoblotting, using the same sample preparation, gel running, transfer and signal development scheme as mentioned before.

*Antibodies used:* ATP5A1 (Proteintech #14676-1-AP), ATP5C1 (Proteintech #10910-1-AP), ATP5B (Proteintech #17247-1-AP), HNRNPC1/C2 (Genetex #14676-1-AP), H3 (Sigma, Upstate #07-690).

#### 2C^2^ Assay

For the first round of 2C, the procedure described above was followed, including on-column DNaseI treatment based on the manufacturer’s protocol (Zymo). Any remaining DNA contamination in the 2C-RNA was then eliminated using TURBO DNAse (Thermo #AM2238) treatment. Samples were treated to either:

(i) DNAse only: 100 µg 2C-RNA was treated with 5 µl TURBO DNAse, for 30 mins at 37 □, 1100 rpm shaking.

(i) DNAse & RNase: 100 µg 2C-RNA was treated with 5 µl TURBO DNAse, 5 µl RNaseI (500 U, Ambion #AM2294) for 30 mins at 37°C, 1100 rpm shaking.

Reactions were stopped by adding 4x volume of Zymo RNA lysis buffer (Zymo #R1060) with subsequent RNA purification as described previously.

*Antibodies used:* ATP5A1 (Rabbit pAb, Proteintech #14676-1-AP; Mouse mAb, abcam #ab14748), ATP5C1 (Proteintech #10910-1-AP), DHX9 (abcam #ab26271), TOM20 (Proteintech #11802-1-AP).

### Polynucleotide Kinase (PNK) assay

PNK assays were based on general principles described before in Horos et al.^3^ with custom optimizations, described below.

#### (i) Antibody-Bead coupling

For each IP set, 20 µl of Protein A SureBead (Bio Rad #161-4811) slurry was equilibrated in PBST-0.05% by three successive washes. Beads were resuspended in 200 µl of PBS with 3 µg of specific antibody and incubated at 4°C with gentle rotation for at least 2 h or overnight. The antibody-bead conjugates were then washed thrice with 500 µl of PBST-0.05% and resuspended in 100 µl of RIPA lysis buffer supplemented with RNase Inhibitor (Ambion) and EDTA-free protease inhibitors (MERCK #04693132001 Roche).

*Antibodies used:* ATP5A1 (Proteintech #14676-1-AP), Rabbit IgG isotype control (CST #2729S).

#### (ii) Preclearing & immunoprecipitation (IP)

For each IP set, 20 µl of Protein A SureBead (Bio Rad #161-4811) slurry was equilibrated to RIPA lysis buffer, by three successive washes. 300 µg of crosslinked Huh7 RIPA lysate was thawed on ice (frozen for maximum 1 month) and added to the beads, 500 µl final volume. Samples were incubated at 4°C with gentle rotation for 2 h.

#### (iii) Nuclease digestion

IP-beads were washed once with a NDB. 100 µl of fresh NDB containing 1U of TURBO DNAse (Thermo #AM2238) and differing amounts of RNase A (Sigma #R5503) was added. Samples were then incubated at 37°C with shaking at 800 rpm for 10 mins. Digestion was stopped by adding 1 µl of RNasIn Plus (Promega #N2611) and chilling on ice for 5 mins.

#### (iv) IP washes

IP-beads were washed (6x) total with 1 ml buffer, 2 mins / wash in following order-(2x) RIPA > (2x) RIPA-HS > (2x) PNK buffer.

#### (v) Radioactive PNK reaction and final washes

IP-beads were resuspended in radioactive PNK reaction mix containing 0.1 µCi/µl [γ-^32^P] ATP (Hartmann analytic #FP-10130), 1 U/µl T4 PNK enzyme (NEB #M0201S) and 5 mM DTT (freshly added). After 15 mins incubation at 37°C, the beads were washed carefully 4x in the PNK buffer. Acidic pH based protein elution (100 mM Glycine–HCl, pH 2.0), neutralisation (200 mM Tris-HCl, pH 8.5), SDS-PAGE and western blotting is described in next section. Phosphorimager exposures were visualized using Typhoon FLA-9500 (GE healthcare) scanner

### Protein co-immunoprecipitation

Cell lysis, chromatin shearing, sample clarification and protein estimation adhered to the previously outlined protocols.Co-immunoprecipitation (Co-IP) assays adapted the general PNK IP scheme with HMGN buffer used consistently for IP and washes. For Flag transgene Co-IPs, different beads were used. Consequently, adjustments were made in specific steps as outlined below:

- Pre-clearing-15 µl of A/G magnetic agarose bead slurry (Pierce # 78609), per 1 mg lysate.
- IP-15 µl Flag-M2 magnetic agarose beads (Sigma # M8823), per 1 mg of lysate.
- Elution-20 µl of elution buffer containing 100-200 µg/ml of Flag peptide (Sigma #F3290) in HMGN, at 4°C for 30 mins. Two eluates were pooled.

### eCLIP library preparation and sequencing

Our eCLIP strategy was based on the one described by Van Nostrad et al. ^5^ previously, with major custom adaptations described below. All buffers were supplemented with EDTA-free protease inhibitors (MERCK #04693132001 Roche) and RNase Inhibitor (Ambion #AM2684), freshly before use.

#### (i) UV crosslinking and lysis

Three UV-crosslinked biological replicates were prepared as described in the previous section. Chromatin was sheared using a Branson tip sonicator and cleared, before snap freezing each replicate set.

#### (ii) Limited nuclease digestion

500 µg lysates (∼300 µl of RIPA) were treated with 1U RNase I (Ambion #AM2294; 100 U/µl stock) and 2U of TURBO DNase (Thermo #AM2238; 2 U/µl stock), for 5 mins at 37°C. Lysates were immediately chilled on ice, centrifuged (10,000 g, 4°C, 10 mins) and supernatants transferred to a new pre-chilled DNA low binding tube, on ice.

#### (iii) Preclearing & IP

For each IP reaction, 500 µg lysate (in 300 µl of RIPA) was used. 1 % lysate was saved for SMI (size matched input). 6 µg of antibodies against either ATP5A1 (Proteintech, rabbit pAb #14676-1-AP; Lot no. 00049350), ATP5C1 (Proteintech, rabbit pAb #10910-1-AP; Lot no. 00029161) or Rabbit IgG isotype control (CST #2729S; Lot no. 08) were used in respective IPs. Bead preparation, antibody coupling, preclearing and IP steps were the same as described for PNK assay. After IP, beads were washed (6x) with 1 ml chilled buffers, 2 mins per wash in following order: (2x) RIPA-HS > (2x) NSWB > (2x) FastAP buffer.

#### (iv) FastAP & PNK treatments of IP beads

Same as mentioned in Supplementary Protocol 1 of {Van Nostrad 2016}.

#### (v) Post-PNK washes of IP beads

Washed 8x total with 0.5 ml chilled buffers, 1 min per wash in following order-(1x) NSWB > (2x) RIPA-HS > (2x) NSWB > (4x) T4 Ligase buffer (1x final concentration).

#### (vi) 3’ Linker ligation reaction of IP beads

Same as mentioned in {PMID: 27018577}, Supplementary Protocol 1.

#### (vii) Post-Linker ligation washes of IP beads

Washed 8x total with 0.5 ml chilled buffers, in rapid succession, with no incubation period. Order-(1x) NSWB > (2x) RIPA-HS > (4x) NSWB.

#### (viii) Elution from IP beads

Elution was carried out using the acidic pH method and then neutralized before gel loading, as previously described for PNK.

#### (ix) Gel electrophoresis and membrane transfer

IP eluates and inputs (saved at beginning of stage iii) were loaded on 4–12% Criterion XT Bis-Tris precast protein gels (Bio-Rad #3450124), run in NuPAGE MOPS SDS running buffer (Invitrogen) supplemented with 1mM DTT, for 75 mins at 150 V. They were transferred to nitrocellulose membranes at 25 V for 30 mins in the Trans-Blot Turbo system (Bio-Rad).

#### (x) Membrane cutting

Individual lanes were cut from the membrane using fresh razor blades. A cutting-mask was placed on top and a consistent size range was applied, spanning just above the target protein band to 75 kDa above it. This approach was maintained across all 3 replicates for both the IP and its corresponding size matched inputs (SMI).

#### (xi) RNA isolation to final library preparation

Same as mentioned in Supplementary Protocol 1 of {Van Nostrad 2016}.

Number of PCR cycles for final cDNA libraries were determined after test PCR runs. Final cycles were determined after selecting

They were: All SMIs (9 cycles); ATP5A1 eCLIP libraries (12 cycles); ATP5C1 eCLIP libraries (10 cycles); Rabbit IgG control eCLIP libraries (14 cycles).

#### (xii) Sequencing

Libraries were multiplexed and subjected to paired-end sequencing (PE 75) using the Illumina NextSeq 500 platform. Illumina i5 (Index1) and i7 (Index2) barcode adaptors were used.

### eCLIP-seq data analysis

The quality of the sequenced eCLIP reads was assessed using FastQC tool (v.0.11.8) ^6^. Unique molecular identifier (UMI) barcodes attached during library preparation were extracted and appended to the read name using UMI-Tools (v1.0.0) ^7^. Adapters were trimmed using Cutadapt tool ^8^ (v2.5) and reads longer than 18 nucleotides were retained.

Next, they were aligned to the human genome (GRCh38.v23 from GENCODE) using STAR (v2.7.1a) ^9^, a splice-aware aligner. Those reads that map to more than 10 genomic locations were discarded using “outFilterMultimapNmax 10” parameter. The other STAR parameters used for mapping were listed as follows: “sjdbOverhang 149”, “outSAMunmapped Within”, “outFilterMultimapScoreRange 1”, “outFilterType BySJout”, “outReadsUnmapped Fastx”, “outFilterScoreMin 10”, “alignEndsType Extend5pOfRead1”, “genomeLoad NoSharedMemory”, “outSAMattributes All”, “outFilterMismatchNoverLmax 0.08”, “seedSearchStartLmax 10”, and the rest are set to default. PCR duplicates were removed from aligned reads using UMI-Tools (dedup command) ^7^, utilizing appended barcodes appended to the read name. The annotation of GRCh38.v23 genome was extended by including coordinates for tRNAs provided by tRNAscan ^10^, and the resulting annotation was pre-processed with the htseq-clip suite ^11^ (v2.0.0) into overlapping windows of 50 nucleotides in size with a step size of 20 nucleotides. The truncation site (position -1 relative to the start site of a read, also called crosslink site) was extracted and quantified using htseq-clip. The R/Bioconductor package DEWSeq ^12^ (v.0.99) was used to detect significantly enriched windows in IP samples over the corresponding size-matched input control samples. Independent hypothesis weighting (IHW) ^13^ was used for multiple hypothesis correction (log_2_ fold change >=1, p_adjusted_ <=0.05). Overlapping significant windows were merged into binding regions with DEWSeq.

### Formaldehyde cross linked RNA Immunoprecipitation (fRIP)

The protocol is detailed in stages.

#### (i) Formaldehyde crosslinking and quenching

Huh7 cells at 90-95% confluency were crosslinked with 0.1% formaldehyde (diluted from a 16% stock in PBS) for 9 minutes at room temperature (RT). After removing the crosslinking solution, cells were washed twice with PBS at RT. Quenching was done with 0.125 M glycine (diluted 20x from a 2.5 M stock in PBS) for 5 minutes with gentle rocking at RT. The glycine was then removed, and cells were washed three times with ice-cold PBS for 1-2 minutes each.

#### (ii) Lysate preparation

The cells were lysed immediately on chilled plates using RIPA lysis buffer (supplied with fresh protease inhibitors and RNase inhibitors), with the same methodology as described before. Frozen lysates were used within 1 month maximum.

#### (iii) Antibody-bead conjugation

20 µl of Protein A magnetic bead slurry (Sure Beads, Bio-Rad) were placed in 1.5 ml DNA low-binding Eppendorf tubes. After buffer removal using a magnet, the beads were equilibrated in 1 ml PBS-T (0.05% Tween 20) and washed three times by inversion. The beads were then incubated with 3 µg antibodies in 300 µl PBS-T, overnight at 4°C with rotation.The antibody-bead conjugate was finally resuspended in 20-25 µl RIPA (for each IP) and stored at 4°C for next day use. Beads for preclearing were prepared in a similar manner with PBS addition instead of antibody. Antibodies used: ATP5A1 (Proteintech, rabbit pAb #14676-1-AP; Lot no. 00049350) or Rabbit IgG isotype control (CST #2729S; Lot no. 08).

#### (iv) Preclearing & IP

For each IP reaction, 500 µg lysate (in 300 µl of RIPA) was used, freshly supplemented with RNase Inhibitor (Ambion) and EDTA-free protease inhibitors (MERCK #04693132001 Roche). Samples were precleared at 4°C, for 30 mins by gentle inversion. 2% lysate was saved for input. IP was performed with anti-ATP5A1 antibody-conjugated beads at 4°C, for 2h by gentle inversion.

#### (v) Post-IP bead washes

Beads were washed 6x total with 1 ml chilled buffers, 2 mins per wash, at RT, in following order-(1x) RIPA > (2x) RIPA-HS > (2x) NSWB > (1x) RIPA.

#### (vi) Proteinase-K treatment & decrosslinking

From this step onwards, both input and IP samples were included. Proteinase-K (PK) reaction mixes were prepared by diluting Proteinase K (20 mg/ml stock, NEB) to a final concentration of 2 mg/ml in the PK buffer and incubated at 37°C for 20 minutes to deactivate any RNAse activity. RNase Inhibitor (Ambion #AM2684) was then added to the PK mix. IP’d beads received 180 µl of the PK reaction buffer, while input samples received 20 µl. They were incubated at 55°C for 30 minutes at 1200 rpm. Afterward, inputs and IP beads were centrifuged, magnetized, and the supernatant (S/N) was retained. Finally, the input volume was adjusted to 180 µl by adding 160 µl of PK reaction buffer, ensuring equal volumes for both IP and input samples in subsequent phases.

#### (vii) Phenol extraction of RNA and purification on silica columns

Input and IP samples were initially treated with 180 µl of acid phenol/chloroform/isoamyl alcohol (pH 6.5) at 37°C, 1200 rpm for 5 minutes to extract RNA. Continuous processing without pauses was used to prevent RNA degradation from prolonged chloroform exposure. Alternatively, Trizol (LS) could be employed for RNA processing at a later stage. After centrifugation at 15,000 rpm for 15 mins at RT, the aqueous layer was transferred to a fresh tube. Subsequent steps utilised the ZYMO RNA Clean & Concentrator-5 kit (Zymo #R1016/1013). RNA purification continued with the addition of RNA binding buffer (double volume) and then equal volume of absolute ethanol. The solution was thoroughly mixed, loaded onto silica columns, and subsequent steps followed the kit instructions.

#### (vii) cDNA preparation and qPCR

All subsequent steps followed respective kit instructions. Purified RNA was treated with 4U TURBO DNase (TURBO DNA-free Kit, Invitrogen #AM1907), supplemented with RNaseOUT RNase Inhibitor (Thermo #10777019). cDNA was prepared from whole or half of purified RNA using SuperScript IV Reverse Transcriptase (Invitrogen #18090200), using random hexamers.

*Primers, human (Fw&Rev):*

RPLPO (TGGTCATCCAGCAGGTGTTCGA & ACAGACACTGGCAACATTGCGG), ATP5C1 (CTCAGACAAGAGGTAAAGAAGG & ACAGTTTCTTCGGACAAAGG), GPC3 (AAAGGACAATCTATATGCTACCACT & GCCATGAAGTAGAGGACTAACCA), CTSD (ATCCCAACCCCACCTCCAGGCCAAT & GGGGACAAAACCCACCTTGTTGGAGCC), RPS20 (GACCAGTTCGAATGCCTACCA & ATCTGGAAACGATCCCACGTC), RPS6 (TGGACGATGAACGCAAACTTC & CGGACCACATAACCCTTCCAT), RPL13 (CTTTCCGCTCGGCTGTTTTC & CCTTGTGGAAGTGGGGCTTC).

### Motif analysis

The genomics intervals of significant binding regions from DEWSeq analysis were converted into respective transcriptomic intervals using a R/Bioconductor package called EnsDb.Hsapiens.v86 and BLAST tool (v2.9.0) ^14^. The converted transcriptomic intervals of protein-coding transcripts expressed in the transcriptome were retained. The intervals were annotated for transcript features, i.e., UTR5, CDS, UTR3, using Bedtools (v.2.27.1) ^15^ and converted into a fasta file.

MEME-ChIP (v5.05) ^16^ from the MEME suite was used to predict de novo motifs. The default, first-order background model was used to generate background sequences from the input query sequences^17^, and the human HOCOMOCO(v11) was used as a motif database^18^. The E-value <= 0.05 was used to filter the significant motifs. Cross-link sites near predicted motif regions were counted to display coverage in 5’UTR and 3’UTR bound regions. Known TOP motif sequences present in the 5’UTR-bound target RNAs were obtained from Cockman et al. ^19^ Using MEME-ChIP, a position-weighted matrix was created. FIMO (v5.3.0) ^20^, with default settings, was then applied to scan these sites, assess eCLIP cross-linking coverage, followed by quantification and visualisation.

### Identification of mitochondrial targeting sequence (MTS)

Human ATP5A1 MTS sequence was identified by combining predictions of two softwares: MitoFates ^21^ - https://mitf.cbrc.pj.aist.go.jp/MitoFates/cgi-bin/top.cgi

MITOPROT II 1.101 ^22^ - https://ihg.gsf.de/ihg/mitoprot.html

Identified N-terminal 43 nucleotide long MTS sequence is: MLSVRVAAA<u>VVRAL</u>PRRAGLVSRNALGSSFIAARNFHASN^#^THL*

*Notable regions are: <u>TOM20 binding site,</u> MPP^#^ and OCT1* cleavage sites*.

It is referred to as wtMTS, mentioned in the figures and hereafter. The key features displayed by the wtMTS were-amphipathic α-helix, presence of TOM20 binding site, OCT1 and MPP recognition/cleavage sites. These features have been reported in multiple mitochondrial precursor proteins that undergo N-terminal MTS cleavage ^23^.

Design of engineered mitochondrial targeting sequence (eMTS): Engineered MTS (eMTS) was derived from the wtMTS of ATP5A1, after proper identification and immunofluorescence based verification.

#### Concept

Our objective was to insert a small tag within the original wtMTS of ATP5A1. This tag would be present in the pre-protein and get cleaved off in the mature protein, resulting in a clear differentiation between the two forms.

#### Important criteria

Inserted tag should:

- Conform with the key mitochondrial targeting and processing features displayed by wtMTS (mentioned before).
- Integrate within the original amino acid sequence of the wtMTS without altering the original MTS conformation (as predicted by Alpha Fold2 ^24^).
- Conform with the key mitochondrial targeting and processing features displayed by wtMTS (mentioned before).
- Not alter the mature protein sequence (based on predicted peptidase cleavage sites and gel migration pattern).

#### Tags tested in silico

We tested multiple tags in silico for their impact on wtMTS targeting and peptidase processing. V5/Myc/HA/Flag tags were inserted at each position, N-terminal to C-terminal, to assess their effect on wtMTS (ATP5A1_11-43_).

Using MitoFates and MITOPROT II 1.101, all 44 possible positions were examined for each tag. This involved assessing mitochondrial targeting scores, along with OCT1 and MPP recognition/cleavage sites. Ultimately, the V5 tag was identified as the optimal candidate for ATP5A1 eMTS design.

The eMTS sequence used is: MLSVRVAAA<u>VVRAL</u>PRRAGLVSRN(GKPIPNPLLGLDST)ALGSSFIAARN^#^FHASNTHL* *Notable regions are: <u>TOM20 binding site,</u> (V5 tag), MPP^#^ and OCT1* cleavage sites*.

### Immunohistochemistry

Conventional immunohistochemistry was performed on Huh7 cells using immunofluorescence based detection strategy. Cells were grown overnight on either glass coverslips or glass bottom multiwell chambers (Ibidi, µ-Slide 18 well #81817) at 50-60 % confluency. Next day, they were fixed at 37°C for 15 mins in a pre-warmed fixation buffer, followed by three PBS washes. Subsequently, they were permeabilized with a permeabilization buffer for 10 mins at room temperature and blocked for 1 h block in an IF buffer. Primary antibodies were effective with at least 1 h incubation at room temperature or overnight (4°C) in the IF buffer.

After three washes in the PBST-0.02% buffer (5 mins each), secondary antibodies or nanobodies were applied in the IF buffer and incubated for 1 h at room temperature. AlexaFluor (647/545/488)-conjugated secondary antibodies against mouse or rabbit IgG were from Molecular Probes (Invitrogen) and equivalent nanobodies were sourced from Chromotek (Proteintech). Following three more washes, 5 mins DAPI (Invitrogen #R37606) staining was performed. Coverslips were curated in ProLong Diamond Antifade Mountant (Invitrogen #P36965) overnight before imaging. For multiwell chambers, liquid mounting medium (Ibidi #5001) was added before imaging.

*Antibodies used:* HA (Biolegend #14676-1-AP), TOM70 (Proteintech #14528-1-AP), FLAG (Sigma #F3165), ATP5A1 (Rabbit pAb, Proteintech #14676-1-AP; Mouse mAb, abcam #ab14748), DIG (abcam #ab76907).

### RNA proximity ligation assays (rnaPLA)

We based our protocol on RNA-protein proximity ligation assays published before ^25,26^. A great detailed adaptation has also been recently published by George et al. ^27^ While the general concept is similar, in our experience, each RBP rnaPLA assay requires adaptations for protein location, protein-RNA affinity, RNA abundance and probe design.

We further developed a new protocol for antibody co-staining and organellar imaging. An optimised protocol for 18-well chambered slide (µ-Slide 18 Well Glass Bottom, Ibidi #81817) is presented, suitable for confocal and superresolution imaging.

#### (i) Cell Seeding

18-well chambered Ibidi slides were coated with Poly-D-lysine (1 mg/ml, Sigma) using ∼50 µl/well for 20 minutes at room temperature. Wells were washed twice with sterile water, air-dried for 5-10 minutes, and stored sterile at 4°C for up to 1 month. For cell seeding, 6000 Huh7 cells were added in 100 µl media per well for next-day fixation.

#### (ii) Fixation

Cells were fixed and in the same manner as used for immunohistochemistry. The chambers were stored in PBS, 4°C for 0-3 days.

#### (iii) Hybridization probe generation

Probe hybridization buffer was warmed to room temperature on a heat block. 100 nM of appropriate probes (Sigma) were added to the hybridization buffer and mixed thoroughly to prepare the hybridization solution. The hybridization solution was then denatured at 60°C for 5 mins and kept at room temperature till use (maximum 10 mins).

#### (iv) Permeabilization & probe hybridization

Cells were permeabilized with a buffer for 10 mins at room temperature before being transferred to a fume hood. Samples were then washed with 100 µl of pre-prepared 0.1 M triethanolamine for 5 mins at room temperature. Acetylation was carried out by adding 3 µl of acetic anhydride per 1 ml of 0.1 M triethanolamine, followed by a 5-min incubation, then adding 6 µl of acetic anhydride per 1 ml of 0.1 M triethanolamine and incubating for an additional 5 mins. These steps aimed to reduce non-specific background. Afterward, samples were washed twice with 100 µl of PBST for 5 mins each at room temperature, followed by two washes with 100 µl of probe hybridization buffer for 2 mins each at room temperature. Subsequently, 50 µl of hybridization solution containing probe was added to each well, denatured by heating at 60°C for 3 mins, and incubated overnight at 37°C in a humidified chamber.

#### (v) Post hybridization washes

All buffers were prepared and wash steps completed in a fume hood. Sequential washes were performed with 200 µl of pre-warmed (37°C) post-hybridization probe wash buffers (a-e).

a. 50 % (v/v) deionized formamide and 5x SSC, at 10 mins, 37°C incubator.
b. 25 % (v/v) deionized formamide and 1x SSC, at 10 mins, 37°C incubator.
c. 12.5 % (v/v) deionized formamide and 2x SSC, at 10 mins, 37°C incubator.
d. 2x SSC and 0.1 % (v/v) Tween 20, at 10 mins, 37°C incubator.
e. 0.2x SSC and 0.1 % (v/v) Tween 20, at 30 mins, 37°C incubator.

In parallel, reconstituted Duolink wash buffers A and B were warmed to room temperature from 4°C.

#### (vi) PLA & Nanobody incubation

Sigma Duolink kit reagents were used, following the kit protocol with custom adaptations where necessary. Primary antibodies against the oligo probe tag (Biotin/DIG), assayed protein (ATP5A1/V5/HA), and co-staining protein (TOM 70/TOM20) were diluted in Duolink antibody diluent and incubated together at 37°C for 1 hour. Subsequent steps, including washes, Duolink PLA (plus + minus) probe antibody incubation, and PLA ligation, remained unchanged.

At the fluorescent reagent amplification stage, the Taq polymerase mixture, following kit based preparation instructions, was spiked with compatible AlexaFluor-labelled secondary nanobodies (from Chromotek, Proteintech). These nanobodies targeted primary antibodies, including the protein assayed in the assayed protein (ATP5A1/V5/HA). The mixture underwent incubation in a pre-heated humidity chamber for 1 hour and 40 minutes at 37°C. This step achieved simultaneous rolling circle signal amplification (RCA), hybridization-based RCA signal detection (fluorescent detection oligo), and nanobody-based protein detection.

#### (vii) Final Washes & Nuclear staining

Samples underwent a series of washes: first with fresh 1x Duolink Wash Buffer B and optional Nuc Blue nuclear stain (Thermo) for 10 mins at room temperature, then with fresh 1x Wash Buffer B for 10 mins at room temperature, followed by a single wash with 0.01x Wash Buffer B (diluted in water) for 2 mins at room temperature. Each well was then covered with Ibidi liquid mounting media (Ibidi) and stored at 4°C before imaging. Imaging was completed within 1-4 days for optimal signal strength, ensuring control and target wells were imaged in the same session.

### rnaPLA Probes, Antibodies & Nanobodies

*3’ Biotin or DIG-labelled antisense probes:*

RPL13 (CTGCCAGTCCTTGTGGAAGTGG), RPS6 (CAGCAACTTCTGTGGCCATACGC), RPS20 (AGAGCGAACAGCGGTGAGTCA), APOE (AGCAACGCAGCCCACAGAACCTT), GPC3 (CTGTAGGGCAGCACATGTGCT).

Primary antibodies used:

ATP5A1 (Rabbit pAb, Proteintech #14676-1-AP; Mouse mAb, abcam #ab14748), DHX9 (abcam #ab26271), TOM20 (Proteintech #11802-1-AP), HA (Mouse mAb, Biolegend #14676-1-AP), TOM70 (Rabbit pAb, Proteintech #14528-1-AP), FLAG-M2 (Mouse mAb, Sigma #F3165), Biotin (Goat pAb, Sigma #B3640; Mouse mAb, abcam #ab201341 ; Rabbit pAb, abcam #ab53494), DIG (Goat pAb, abcam #ab76907), V5 (Mouse mAb,Thermo #R960-25).

AlexaFluor conjugated Nanobodies (Proteintech):

Anti-IgG Rb Nanobody (clones-CTK0101, CTK0102) #srbAF488,568,647, anti IgG2b Ms Nanobody #sms2bAF488.

### Image acquisition

#### Confocal microscopy

Standard confocal images were captured with Olympus FV3000, using PLAPON 60X OSC /NA 1.4 / Oil WD 0.12 mm or UPLSAPO 60X S / NA 1.3 / Silicon WD 0.3 mm objectives, in Z-stacks. It featured galvano and resonant scanning, four spectral GaAsP detectors, and Olympus IX3-ZDC2 Z Drift Compensation.

#### Super Resolution confocal microscopy

Superresolution images were captured using the Zeiss LSM880 AiryFast microscope in Airyscan SR imaging mode, using methodologies mentioned before^28^. Images were obtained by employing a main beam splitter (MBS 488/561/633) for excitation and standard filters for detection. The imaging was carried out using either the Plan-Apochromat 40x/1.4 Oil DIC M27 (FWD=0.13mm) or Plan-Apochromat 63x/1.4 Oil DIC M27 objectives. Subsequent image processing, including deconvolution and pixel reassignment, was conducted in Zen Black software (Carl Zeiss). We used either 2D (single plane) or 3D (Z-Stacks) parameters for deconvolution, depending on the acquired images; before further analysis.

### Image analysis

Images underwent post-processing in Fiji software (V2.1.0) ^29^ (https://imagej.net/software/fiji/) and were converted to TIFF format. 3D (Z-stacks) images were converted to single plane images, though maximum intensity projection beforehand. They were analysed using custom scripts in CellProfiler (V4.06) ^30^ software (http://cellprofiler.org).

### Mitochondria isolation and *in vitro* import assays

Mitochondria were isolated from HEK293T cells by subcellular fractionation ^31^. Cells were rinsed with PBS, resuspended in 83 mM sucrose, 10 mM HEPES pH 7.2 with 1 mM PMSF, and homogenised with a glass-teflon homogeniser. After addition of an equal volume of 250 mM sucrose, 30 mM HEPES pH 7.2 with 1 mM PMSF, the cell lysates were cleared by centrifugation at 1000 g and the mitochondrial fraction was pelleted at 10,000 g. The mitochondria were resuspended in 320 mM sucrose, 1 mM EDTA, 10 mM Tris-HCl pH 7.4 with 1 mM PMSF. RNase A (Thermo Scientific #EN0531) was added to a concentration of 10 ng/µl or omitted and the samples were incubated at 30°C for 10 mins. Immediately afterwards, murine RNase inhibitor (NEB #M0314L) was added to a final concentration of 0.25-1 U/µl and the samples were incubated on ice for 10 mins. Mitochondria were re-isolated and resuspended in import buffer (3 % BSA, 250 mM sucrose, 5 mM magnesium acetate, 80 mM potassium acetate, 10 mM sodium succinate, 1 mM DTT, 5 mM ATP, 20 mM HEPES pH 7.4) ^32^, supplemented with 0.25-0.5 U/µl murine RNase inhibitor. Radiolabeled ATP5A1 precursor protein was generated in vitro as follows. The ORF was amplified using the template ATP5A1 cDNA template and primers SP6-EcoRI-ATP5A1-F (tcgatttaggtgacactatagaagtgaattcatgctgtccgtgagagttg) or SP6-ATP5A1-F (gatcgatttaggtgacactatagaagcggccaccatgctgtccgtgagagttg) and ATP5A1-R (ctatgtctagattaagcttcaaatccagccaag). Transcription and translation in the presence of [^35^S] methionine were performed with the mMessage mMachine kit (Ambion #AM1340) and Flexi Rabbit Reticulocyte Lysate (Promega #L4540), respectively. Radiolabeled rat ornithine transcarbamylase (OTC) precursor protein was synthesized from plasmid OTC with the TNT Quick Coupled Transcription/Translation kit (Promega #L2080). Upon import into the mitochondrial matrix, the OTC precursor is subject to two sequential processing steps by MPP (mitochondrial processing peptidase) and MIP (mitochondrial intermediate peptidase) ^33,34^ ^35^. Import of radiolabeled precursors was performed at 30°C. The mitochondrial membrane potential (Δψ) was dissipated by addition of 1 µM valinomycin to stop import or, for the -Δψ control samples, prior to import. Proteinase K treatment was performed with 50 µg/ml PK on ice followed by PMSF addition. Mitochondria were pelleted, washed with 250 mM sucrose, 1 mM EDTA, 10 mM MOPS pH 7.2, and the samples were analysed by SDS-PAGE and autoradiography.

### Electrophoretic mobility shift assay (EMSA) experiments

EMSA binding buffer (10 mM Tris, 150 mM NaCl, 0.5 mM DTT, 0.01 µg/µl BSA, 5 mM MgCl_2_) was used for dilution of protein and RNA components, as well as RNA-Protein (RNP) complex formation. 10 nM fixed concentration of RNA ligand (Cy5 labelled at 3’end, HPLC purified, IDT) was incubated with increasing titrations of protein concentration. The EMSA reaction was incubated at 25°C for 20 mins. Resultant RNA-Protein (RNP) complexes were resolved by native polyacrylamide gel electrophoresis (4-15% Tris-Glycine gradient gels, Bio-Rad #5671084) at 4°C, with a pre-chilled running buffer (34 mM Tris-HCl, 66 mM HEPES, 0.1 mM EDTA) at 100 V for 75 mins. The fluorescent signals were visualized using the Typhoon FLA-9500 (GE healthcare) scanner with appropriate filters.

*RNA fragments:*

RPL13 RNA ligand-CUUUCCGCUCGGCUGUUUUCCUGCGCAGGAGCCGCA scrRPL13 RNA ligand-UCUCUCCGUUCUGCUCGUCGCUGCGCAGAGAGUCGC

## Main References

1. Busch, J. D., Fielden, L. F., Pfanner, N. & Wiedemann, N. Mitochondrial protein transport: Versatility of translocases and mechanisms. Mol. Cell 83, 890–910 (2023).

2. Becker, T., Song, J. & Pfanner, N. Versatility of Preprotein Transfer from the Cytosol to Mitochondria. Trends Cell Biol. 29, 534–548 (2019).

3. Palmer, C. S., Anderson, A. J. & Stojanovski, D. Mitochondrial protein import dysfunction: mitochondrial disease, neurodegenerative disease and cancer. FEBS Lett. 595, 1107–1131 (2021).

4. Den Brave, F., Schulte, U., Fakler, B., Pfanner, N. & Becker, T. Mitochondrial complexome and import network. Trends Cell Biol. (2023) doi:10.1016/j.tcb.2023.10.004.

5. Tsvetanova, N. G., Klass, D. M., Salzman, J. & Brown, P. O. Proteome-wide search reveals unexpected RNA-binding proteins in Saccharomyces cerevisiae. PLoS One 5, (2010).

6. Matia-González, A. M., Laing, E. E. & Gerber, A. P. Conserved mRNA-binding proteomes in eukaryotic organisms. Nat. Struct. Mol. Biol. 22, 1027–1033 (2015).

7. Asencio, C., Chatterjee, A. & Hentze, M. W. Silica-based solid-phase extraction of cross- linked nucleic acid-bound proteins. Life Sci Alliance 1, e201800088 (2018).

8. Wessels, H.-H. et al. The mRNA-bound proteome of the early fly embryo. Genome Res. 26, 1000–1009 (2016).

9. Panhale, A. et al. CAPRI enables comparison of evolutionarily conserved RNA interacting regions. Nat. Commun. 10, 2682 (2019).

10. Marondedze, C., Thomas, L., Serrano, N. L., Lilley, K. S. & Gehring, C. The RNA-binding protein repertoire of Arabidopsis thaliana. Sci. Rep. 6, 29766 (2016).

11. Marondedze, C., Thomas, L., Gehring, C. & Lilley, K. S. Changes in the Arabidopsis RNA- binding proteome reveal novel stress response mechanisms. BMC Plant Biol. 19, 139 (2019).

12. Zhang, Z. et al. UV crosslinked mRNA-binding proteins captured from leaf mesophyll protoplasts. Plant Methods 12, 42 (2016).

13. Boucas, J. et al. Label-Free Protein-RNA Interactome Analysis Identifies Khsrp Signaling Downstream of the p38/Mk2 Kinase Complex as a Critical Modulator of Cell Cycle Progression. PLoS One 10, e0125745 (2015).

14. He, C. et al. High-Resolution Mapping of RNA-Binding Regions in the Nuclear Proteome of Embryonic Stem Cells. Mol. Cell 64, 416–430 (2016).

15. Liao, Y. et al. The Cardiomyocyte RNA-Binding Proteome: Links to Intermediary Metabolism and Heart Disease. Cell Rep. 16, 1456–1469 (2016).

16. Liepelt, A. et al. Identification of RNA-binding Proteins in Macrophages by Interactome Capture. Mol. Cell. Proteomics 15, 2699–2714 (2016).

17. Perez-Perri, J. I. et al. The RNA-binding protein landscapes differ between mammalian organs and cultured cells. Nat. Commun. 14, 2074 (2023).

18. Castello, A. et al. Insights into RNA biology from an atlas of mammalian mRNA-binding proteins. Cell 149, 1393–1406 (2012).

19. Beckmann, B. M. et al. The RNA-binding proteomes from yeast to man harbour conserved enigmRBPs. Nat. Commun. 6, 10127 (2015).

20. Castello, A. et al. Comprehensive Identification of RNA-Binding Domains in Human Cells. Mol. Cell 63, 696–710 (2016).

21. Horos, R. et al. The Small Non-coding Vault RNA1-1 Acts as a Riboregulator of Autophagy. Cell 176, 1054–1067.e12 (2019).

22. Caudron-Herger, M. et al. R-DeeP: Proteome-wide and Quantitative Identification of RNA- Dependent Proteins by Density Gradient Ultracentrifugation. Mol. Cell 75, 184–199.e10 (2019).

23. Mullari, M., Lyon, D., Jensen, L. J. & Nielsen, M. L. Specifying RNA-Binding Regions in Proteins by Peptide Cross-Linking and Affinity Purification. J. Proteome Res. 16, 2762–2772 (2017).

24. Trendel, J. et al. The Human RNA-Binding Proteome and Its Dynamics during Translational Arrest. Cell 176, 391–403.e19 (2019).

25. Teixeira, F. K. et al. ATP synthase promotes germ cell differentiation independent of oxidative phosphorylation. Nat. Cell Biol. 17, 689–696 (2015).

26. Shin, B. et al. Mitochondrial Oxidative Phosphorylation Regulates the Fate Decision between Pathogenic Th17 and Regulatory T Cells. Cell Rep. 30, 1898–1909.e4 (2020).

27. Galber, C., Carissimi, S., Baracca, A. & Giorgio, V. The ATP Synthase Deficiency in Human Diseases. Life 11, (2021).

28. Galber, C., Acosta, M. J., Minervini, G. & Giorgio, V. The role of mitochondrial ATP synthase in cancer. Biol. Chem. 401, 1199–1214 (2020).

29. Jonckheere, A. I., Smeitink, J. A. M. & Rodenburg, R. J. T. Mitochondrial ATP synthase: architecture, function and pathology. J. Inherit. Metab. Dis. 35, 211–225 (2012).

30. Van Nostrand, E. L. et al. Robust transcriptome-wide discovery of RNA-binding protein binding sites with enhanced CLIP (eCLIP). Nat. Methods 13, 508–514 (2016).

31. Sahadevan, S., Sekaran, T. & Schwarzl, T. A Pipeline for Analyzing eCLIP and iCLIP Data with Htseq-clip and DEWSeq. Methods Mol. Biol. 2404, 189–205 (2022).

32. Cockman, E., Anderson, P. & Ivanov, P. TOP mRNPs: Molecular Mechanisms and Principles of Regulation. Biomolecules 10, (2020).

33. Schäfer, J. A., Bozkurt, S., Michaelis, J. B., Klann, K. & Münch, C. Global mitochondrial protein import proteomics reveal distinct regulation by translation and translocation machinery. Mol. Cell 82, 435–446.e7 (2022).

34. Pfanner, N. Protein sorting: recognizing mitochondrial presequences. Curr. Biol. 10, R412–5 (2000).

35. Poveda-Huertes, D., Mulica, P. & Vögtle, F. N. The versatility of the mitochondrial presequence processing machinery: cleavage, quality control and turnover. Cell Tissue Res. 367, 73–81 (2017).

36. Castello, A. et al. Identification of RNA-binding domains of RNA-binding proteins in cultured cells on a system-wide scale with RBDmap. Nat. Protoc. 12, 2447–2464 (2017).

37. Fuller, G. G. et al. RNA promotes phase separation of glycolysis enzymes into yeast G bodies in hypoxia. Elife 9, (2020).

38. Brandina, I. et al. Enolase takes part in a macromolecular complex associated to mitochondria in yeast. Biochim. Biophys. Acta 1757, 1217–1228 (2006).

39. Dollenmaier, G. & Weitz, M. Interaction of glyceraldehyde-3-phosphate dehydrogenase with secondary and tertiary RNA structural elements of the hepatitis A virus 3’ translated and non- translated regions. J. Gen. Virol. 84, 403–414 (2003).

40. Singh, R. & Green, M. R. Sequence-specific binding of transfer RNA by glyceraldehyde-3- phosphate dehydrogenase. Science 259, 365–368 (1993).

41. Nagy, E. et al. Identification of the NAD(+)-binding fold of glyceraldehyde-3-phosphate dehydrogenase as a novel RNA-binding domain. Biochem. Biophys. Res. Commun. 275, 253–260 (2000).

42. Zhou, Y. et al. The multifunctional protein glyceraldehyde-3-phosphate dehydrogenase is both regulated and controls colony-stimulating factor-1 messenger RNA stability in ovarian cancer. Mol. Cancer Res. 6, 1375–1384 (2008).

43. Huppertz, I. et al. Riboregulation of Enolase 1 activity controls glycolysis and embryonic stem cell differentiation. Mol. Cell 82, 2666–2680.e11 (2022).

44. Rabinovitz, M. Uncharged tRNA-phosphofructokinase interaction in amino acid deficiency. Amino Acids 10, 99–108 (1996).

45. Scherrer, T., Mittal, N., Janga, S. C. & Gerber, A. P. A screen for RNA-binding proteins in yeast indicates dual functions for many enzymes. PLoS One 5, e15499 (2010).

46. Kiri, A. & Goldspink, G. RNA-protein interactions of the 3’ untranslated regions of myosin heavy chain transcripts. J. Muscle Res. Cell Motil. 23, 119–129 (2002).

47. Shen, T., Wang, H., Tang, B., Zhu, G. & Wang, X. The impact of RNA binding proteins and the associated long non-coding RNAs in the TCA cycle on cancer pathogenesis. RNA Biol. 20, 223–234 (2023).

48. Chen, Y.-H. et al. MDH2 is an RNA binding protein involved in downregulation of sodium channel Scn1a expression under seizure condition. Biochim. Biophys. Acta Mol. Basis Dis. 1863, 1492–1499 (2017).

49. Xu, F. et al. LncRNA AC020978 facilitates non-small cell lung cancer progression by interacting with malate dehydrogenase 2 and activating the AKT pathway. Cancer Sci. 112, 4501–4514 (2021).

50. Sang, L. et al. Mitochondrial long non-coding RNA GAS5 tunes TCA metabolism in response to nutrient stress. Nat Metab 3, 90–106 (2021).

51. Noble, M., Chatterjee, A., Sekaran, T., Schwarzl, T. & Hentze, M. W. Cytosolic RNA binding of the mitochondrial TCA cycle enzyme malate dehydrogenase (MDH2). RNA (2024) doi:10.1261/rna.079925.123.

52. Xiang, S. et al. LncRNA IDH1-AS1 links the functions of c-Myc and HIF1α via IDH1 to regulate the Warburg effect. Proc. Natl. Acad. Sci. U. S. A. 115, E1465–E1474 (2018).

53. Zhao, H. et al. The opposite role of alternatively spliced isoforms of LINC00477 in gastric cancer. Cancer Manag. Res. 11, 4569–4576 (2019).

54. Zheng, Z.-Q. et al. Long Noncoding RNA TINCR-Mediated Regulation of Acetyl-CoA Metabolism Promotes Nasopharyngeal Carcinoma Progression and Chemoresistance. Cancer Res. 80, 5174–5188 (2020).

55. Hwang, H. J. & Kim, Y. K. The role of LC3B in autophagy as an RNA-binding protein. Autophagy 19, 1028–1030 (2023).

## Supplementary References

1. Noble, M., Chatterjee, A., Sekaran, T., Schwarzl, T. & Hentze, M. W. Cytosolic RNA binding of the mitochondrial TCA cycle enzyme malate dehydrogenase (MDH2). RNA (2024) doi:10.1261/rna.079925.123.

2. Chatterjee, A. et al. MOF Acetyl Transferase Regulates Transcription and Respiration in Mitochondria. Cell 167, 722–738.e23 (2016).

3. Horos, R. et al. The Small Non-coding Vault RNA1-1 Acts as a Riboregulator of Autophagy. Cell 176, 1054–1067.e12 (2019).

4. Asencio, C., Chatterjee, A. & Hentze, M. W. Silica-based solid-phase extraction of cross-linked nucleic acid-bound proteins. Life Sci Alliance 1, e201800088 (2018).

5. Van Nostrand, E. L. et al. Robust transcriptome-wide discovery of RNA-binding protein binding sites with enhanced CLIP (eCLIP). Nat. Methods 13, 508–514 (2016).

6. Andrews, S. & Krueger, F. FastQC. quality control tool … (2010).

7. Smith, T., Heger, A. & Sudbery, I. UMI-tools: modeling sequencing errors in Unique Molecular Identifiers to improve quantification accuracy. Genome Res. 27, 491–499 (2017).

8. Martin, M. Cutadapt removes adapter sequences from high-throughput sequencing reads. EMBnet.journal 17, 10–12 (2011).

9. Dobin, A. et al. STAR: ultrafast universal RNA-seq aligner. Bioinformatics 29, 15–21 (2013).

10. Lowe, T. M. & Chan, P. P. tRNAscan-SE On-line: integrating search and context for analysis of transfer RNA genes. Nucleic Acids Res. 44, W54–7 (2016).

11. Sahadevan, S. et al. htseq-clip: a toolset for the preprocessing of eCLIP/iCLIP datasets. Bioinformatics 39, (2023).

12. Sahadevan, S., Sekaran, T. & Schwarzl, T. A Pipeline for Analyzing eCLIP and iCLIP Data with Htseq-clip and DEWSeq. Methods Mol. Biol. 2404, 189–205 (2022).

13. Ignatiadis, N., Klaus, B., Zaugg, J. B. & Huber, W. Data-driven hypothesis weighting increases detection power in genome-scale multiple testing. Nat. Methods 13, 577–580 (2016).

14. Mount, D. W. Using the Basic Local Alignment Search Tool (BLAST). CSH Protoc. 2007, db.top17 (2007).

15. Quinlan, A. R. & Hall, I. M. BEDTools: a flexible suite of utilities for comparing genomic features. Bioinformatics 26, 841–842 (2010).

16. Machanick, P. & Bailey, T. L. MEME-ChIP: motif analysis of large DNA datasets. Bioinformatics 27, 1696–1697 (2011).

17. Ma, W., Noble, W. S. & Bailey, T. L. Motif-based analysis of large nucleotide data sets using MEME-ChIP. Nat. Protoc. 9, 1428–1450 (2014).

18. Kulakovskiy, I. V. et al. HOCOMOCO: a comprehensive collection of human transcription factor binding sites models. Nucleic Acids Res. 41, D195–202 (2013).

19. Cockman, E., Anderson, P. & Ivanov, P. TOP mRNPs: Molecular Mechanisms and Principles of Regulation. Biomolecules 10, (2020).

20. Grant, C. E., Bailey, T. L. & Noble, W. S. FIMO: scanning for occurrences of a given motif. Bioinformatics 27, 1017–1018 (2011).

21. Fukasawa, Y. et al. MitoFates: improved prediction of mitochondrial targeting sequences and their cleavage sites. Mol. Cell. Proteomics 14, 1113–1126 (2015).

22. Claros, M. G. & Vincens, P. Computational method to predict mitochondrially imported proteins and their targeting sequences. Eur. J. Biochem. 241, 779–786 (1996).

23. Wiedemann, N. & Pfanner, N. Mitochondrial Machineries for Protein Import and Assembly. Annu. Rev. Biochem. 86, 685–714 (2017).

24. Jumper, J. et al. Highly accurate protein structure prediction with AlphaFold. Nature 596, 583–589 (2021).

25. Jung, J., Lifland, A. W., Alonas, E. J., Zurla, C. & Santangelo, P. J. Characterization of mRNA-cytoskeleton interactions in situ using FMTRIP and proximity ligation. PLoS One 8, e74598 (2013).

26. Zhang, W., Xie, M., Shu, M.-D., Steitz, J. A. & DiMaio, D. A proximity-dependent assay for specific RNA-protein interactions in intact cells. RNA 22, 1785–1792 (2016).

27. George, J., Mittal, S., Kadamberi, I. P., Pradeep, S. & Chaluvally-Raghavan, P. Optimized proximity ligation assay (PLA) for detection of RNA-protein complex interactions in cell lines. STAR Protoc 3, 101340 (2022).

28. Wu, X. & Hammer, J. A. ZEISS Airyscan: Optimizing Usage for Fast, Gentle, Super-Resolution Imaging. Methods Mol. Biol. 2304, 111–130 (2021).

29. Schindelin, J., et al. Fiji: an open-source platform for biological-image analysis. Nat. Methods 9, 676–682 (2012).

30. Stirling, D. R. et al. CellProfiler 4: improvements in speed, utility and usability. BMC Bioinformatics 22, 433 (2021).

31. Acín-Pérez, R., Fernández-Silva, P., Peleato, M. L., Pérez-Martos, A. & Enriquez, J. A. Respiratory active mitochondrial supercomplexes. Mol. Cell 32, 529–539 (2008).

32. Kang, Y. et al. Sengers Syndrome-Associated Mitochondrial Acylglycerol Kinase Is a Subunit of the Human TIM22 Protein Import Complex. Mol. Cell 67, 457–470.e5 (2017).

33. Sztul, E. S. et al. Import of rat ornithine transcarbamylase precursor into mitochondria: two-step processing of the leader peptide. J. Cell Biol. 105, 2631–2639 (1987).

34. Kalousek, F., Hendrick, J. P. & Rosenberg, L. E. Two mitochondrial matrix proteases act sequentially in the processing of mammalian matrix enzymes. Proc. Natl. Acad. Sci. U. S. A. 85, 7536–7540 (1988).

