## Supplementary data table 1 for "RNA promotes mitochondrial import of F1-ATP synthase subunit alpha (ATP5A1)"

### Extended Data Table 1

| Gene Id | Gene Names | Gene Types | ATP5A1-IP_Average | ATP5A1-SMI_Average |
|---|---|---|---|---|
| ENSG00000149930.17 | TAOK2 | protein_coding | 25.00486382 | 5.218198008 |
| ENSG00000222923.1 | RNU2-41P | snRNA | 25.13627117 | 3.902982463 |
| ENSG00000196683.10 | TOMM7 | protein_coding | 25.2493734 | 8.512167167 |
| ENSG00000159335.15 | PTMS | protein_coding | 25.37496431 | 7.136002154 |
| ENSG00000199420.1 | RN7SKP170 | misc_RNA | 25.76226811 | 2.113445071 |
| ENSG00000160803.7 | UBQLN4 | protein_coding | 25.76774321 | 7.036250631 |
| ENSG00000182087.12 | TMEM259 | protein_coding | 25.91175655 | 7.126968179 |
| ENSG00000223668.1 | EEF1A1P24 | processed_pseudogene | 25.96605368 | 2.021474674 |
| ENSG00000143321.18 | HDGF | protein_coding | 26.25590871 | 9.509899738 |
| ENSG00000080824.18 | HSP90AA1 | protein_coding | 26.29009053 | 8.935826369 |
| ENSG00000219200.10 | RNASEK | protein_coding | 26.47422778 | 9.368680004 |
| ENSG00000161939.18 | RNASEK-C17orf49 | protein_coding | 26.47422778 | 9.368680004 |
| ENSG00000128309.16 | MPST | protein_coding | 26.52190632 | 7.479391089 |
| ENSG00000168802.12 | CHTF8 | protein_coding | 26.58710676 | 7.428888903 |
| ENSG00000198952.8 | SMG5 | protein_coding | 26.66497146 | 7.498872346 |
| ENSG00000127837.9 | AAMP | protein_coding | 26.98606757 | 6.201724749 |
| ENSG00000213235.3 | EEF1A1P16 | processed_pseudogene | 26.98745245 | 1.1157125 |
| ENSG00000222259.1 | RN7SKP114 | misc_RNA | 27.02967326 | 0.743808334 |
| ENSG00000126432.13 | PRDX5 | protein_coding | 27.10743281 | 10.4843925 |
| 20281 | 20281 | tRNAscan | 27.65221637 | 7.028469506 |
| ENSG00000197746.13 | PSAP | protein_coding | 27.73199719 | 12.17543122 |
| ENSG00000151176.7 | PLBD2 | protein_coding | 27.91931657 | 10.83681541 |
| ENSG00000134824.13 | FADS2 | protein_coding | 28.0032754 | 10.39492781 |
| ENSG00000211459.2 | MT-RNR1 | Mt_rRNA | 28.07726175 | 9.87276993 |
| ENSG00000132613.14 | MTSS1L | protein_coding | 28.20350199 | 8.223199423 |
| ENSG00000175634.14 | RPS6KB2 | protein_coding | 28.61615325 | 8.78557266 |
| ENSG00000064666.14 | CNN2 | protein_coding | 28.66434488 | 10.9975164 |
| ENSG00000198911.11 | SREBF2 | protein_coding | 28.94230072 | 10.07227298 |
| ENSG00000114867.19 | EIF4G1 | protein_coding | 28.96651684 | 12.43196473 |
| ENSG00000211450.9 | C11orf31 | protein_coding | 28.99575401 | 11.58062375 |
| ENSG00000155366.16 | RHOC | protein_coding | 29.11797967 | 7.620610823 |
| ENSG00000068400.13 | GRIPAP1 | protein_coding | 29.44043192 | 7.003816394 |
| ENSG00000171067.10 | C11orf24 | protein_coding | 29.48152139 | 9.648613773 |
| ENSG00000158874.11 | APOA2 | protein_coding | 29.54056945 | 10.54659482 |
| ENSG00000101160.13 | CTSZ | protein_coding | 29.64302712 | 8.764838555 |
| ENSG00000171552.12 | BCL2L1 | protein_coding | 29.71525174 | 13.60209842 |
| ENSG00000117984.12 | CTSD | protein_coding | 30.01175488 | 11.29040315 |
| ENSG00000250644.2 | RP11-295K3.1 | protein_coding | 30.01175488 | 11.29040315 |
| ENSG00000189060.5 | H1F0 | protein_coding | 30.08571249 | 7.16843639 |
| ENSG00000130706.12 | ADRM1 | protein_coding | 30.1154506 | 8.133734724 |
| ENSG00000167685.14 | ZNF444 | protein_coding | 30.1336029 | 2.049989905 |
| ENSG00000152926.14 | ZNF117 | protein_coding | 30.29548547 | 2.71728652 |
| ENSG00000068308.13 | OTUD5 | protein_coding | 30.32113493 | 12.24541466 |
| ENSG00000092841.18 | MYL6 | protein_coding | 30.47161925 | 11.89157845 |
| ENSG00000258643.5 | BCL2L2-PABPN1 | protein_coding | 30.47424218 | 7.99251499 |
| ENSG00000141522.11 | ARHGDIA | protein_coding | 30.52151357 | 9.559149074 |
| ENSG00000198931.10 | APRT | protein_coding | 30.61914547 | 9.41792934 |
| ENSG00000143420.17 | ENSA | protein_coding | 30.77932614 | 11.67912242 |
| ENSG00000171552.12 | BCL2L1 | protein_coding | 31.22089598 | 8.313916971 |

| | | | | |
|---|---|---|---|---|
| ENSG00000117984.12 | CTSD | protein_coding | 31.28761676 | 9.68104801 |
| ENSG00000132471.11 | WBP2 | protein_coding | 31.3245759 | 7.782564663 |
| ENSG00000178952.8 | TUFM | protein_coding | 31.41466975 | 5.408667079 |
| ENSG00000183255.11 | PTTG1IP | protein_coding | 31.41631339 | 10.31340469 |
| ENSG00000248919.7 | ATP5J2-PTCD1 | protein_coding | 31.49063648 | 11.15947024 |
| ENSG00000011304.16 | PTBP1 | protein_coding | 31.57628141 | 15.05728085 |
| ENSG00000199731.1 | RNU6-1079P | snRNA | 31.79990981 | 5.911504156 |
| ENSG00000182747.4 | SLC35D3 | protein_coding | 31.90795008 | 0.603841449 |
| ENSG00000167685.14 | ZNF444 | protein_coding | 32.01450492 | 9.178210933 |
| ENSG00000114391.12 | RPL24 | protein_coding | 32.10764544 | 13.10970863 |
| ENSG00000257529.5 | RPL36A-HNRNPH2 | protein_coding | 32.24024395 | 13.43236346 |
| ENSG00000252623.1 | RNA5SP481 | rRNA | 32.88561751 | 3.608842864 |
| ENSG00000241343.9 | RPL36A | protein_coding | 33.07648008 | 13.43236346 |
| ENSG00000197694.13 | SPTAN1 | protein_coding | 33.08209063 | 15.28143701 |
| ENSG00000100836.10 | PABPN1 | protein_coding | 33.31733073 | 9.74183702 |
| ENSG00000202198.1 | RN7SK | misc_RNA | 33.46624104 | 9.499612914 |
| ENSG00000173020.10 | ADRBK1 | protein_coding | 33.49263283 | 15.23218767 |
| ENSG00000202392.1 | RN7SKP292 | misc_RNA | 33.71093296 | 3.055503601 |
| ENSG00000087460.23 | GNAS | protein_coding | 33.86959329 | 10.1759435 |
| 28108 | 28108 | tRNAscan | 33.97653097 | 3.629576969 |
| ENSG00000169710.6 | FASN | protein_coding | 34.18499106 | 12.94634148 |
| ENSG00000134287.9 | ARF3 | protein_coding | 34.28621146 | 11.55085567 |
| ENSG00000272822.1 | RP11-302B13.5 | protein_coding | 34.28621146 | 11.55085567 |
| ENSG00000201793.1 | RN7SKP9 | misc_RNA | 34.30201175 | 2.372644735 |
| ENSG00000047849.21 | MAP4 | protein_coding | 34.37690697 | 8.322950946 |
| ENSG00000166681.13 | NGFRAP1 | protein_coding | 34.49088776 | 8.153215981 |
| ENSG00000120885.19 | CLU | protein_coding | 34.70426219 | 14.81756244 |
| ENSG00000201583.1 | RN7SKP191 | misc_RNA | 35.17701485 | 0.905762173 |
| ENSG00000126247.10 | CAPNS1 | protein_coding | 35.4227814 | 12.37368142 |
| ENSG00000200779.1 | RNU6-105P | snRNA | 35.44408631 | 4.565107223 |
| ENSG00000007520.3 | TSR3 | protein_coding | 35.49473511 | 8.843855972 |
| ENSG00000168488.18 | ATXN2L | protein_coding | 35.5041885 | 9.000694843 |
| ENSG00000130175.9 | PRKCSH | protein_coding | 36.12207603 | 9.127708747 |
| ENSG00000067182.7 | TNFRSF1A | protein_coding | 36.12904195 | 8.394187237 |
| ENSG00000242372.6 | EIF6 | protein_coding | 36.27359791 | 12.86607121 |
| ENSG00000261582.1 | RP4-614O4.11 | antisense | 36.27359791 | 12.86607121 |
| ENSG00000107223.12 | EDF1 | protein_coding | 36.31102169 | 12.11448175 |
| ENSG00000134824.13 | FADS2 | protein_coding | 36.35931925 | 17.24974334 |
| ENSG00000075618.17 | FSCN1 | protein_coding | 36.58281298 | 10.72536376 |
| ENSG00000168758.10 | SEMA4C | protein_coding | 37.60131869 | 5.871288794 |
| ENSG00000172270.18 | BSG | protein_coding | 37.62678345 | 16.30643196 |
| ENSG00000258674.5 | CTC-260F20.3 | protein_coding | 37.79422969 | 16.41271176 |
| ENSG00000174684.6 | B4GAT1 | protein_coding | 37.8030058 | 15.92424097 |
| ENSG00000173762.7 | CD7 | protein_coding | 37.96594329 | 13.21724127 |
| ENSG00000188157.13 | AGRN | protein_coding | 38.18055498 | 13.70963107 |
| ENSG00000168237.17 | GLYCTK | protein_coding | 38.68747965 | 10.44417714 |
| ENSG00000087088.19 | BAX | protein_coding | 38.72800993 | 17.62164751 |
| ENSG00000167244.17 | IGF2 | protein_coding | 39.03468985 | 12.52393513 |
| ENSG00000197136.4 | PCNXL3 | protein_coding | 39.19398059 | 7.504044201 |
| ENSG00000159720.11 | ATP6V0D1 | protein_coding | 39.22490308 | 11.47836653 |
| ENSG00000214753.2 | HNRNPUL2 | protein_coding | 39.30632754 | 10.51290774 |
| ENSG00000149257.13 | SERPINH1 | protein_coding | 39.52609732 | 11.89935957 |

| | | | | |
|---|---|---|---|---|
| ENSG00000258674.5 | CTC-260F20.3 | protein_coding | 39.74244121 | 18.12573743 |
| ENSG00000116133.11 | DHCR24 | protein_coding | 40.25045033 | 17.37022897 |
| ENSG00000250644.2 | RP11-295K3.1 | protein_coding | 40.25979779 | 19.58358602 |
| ENSG00000135390.17 | ATP5G2 | protein_coding | 40.94624424 | 19.65607516 |
| ENSG00000201517.1 | RNU6-707P | snRNA | 40.97979645 | 3.98074703 |
| ENSG00000101412.12 | E2F1 | protein_coding | 41.14142625 | 3.778577829 |
| ENSG00000119383.19 | PPP2R4 | protein_coding | 41.15923044 | 14.47542636 |
| ENSG00000068028.17 | RASSF1 | protein_coding | 41.30499415 | 8.396853394 |
| ENSG00000089597.16 | GANAB | protein_coding | 41.60825055 | 14.31472537 |
| ENSG00000177733.6 | HNRNPA0 | protein_coding | 41.67892688 | 10.33272549 |
| ENSG00000281621.1 | Y_RNA | misc_RNA | 42.0522496 | 4.324135966 |
| ENSG00000071082.10 | RPL31 | protein_coding | 42.08705123 | 18.30466683 |
| ENSG00000132507.17 | EIF5A | protein_coding | 42.09587344 | 16.72491931 |
| ENSG00000170266.15 | GLB1 | protein_coding | 42.14350509 | 15.56137078 |
| ENSG00000160783.19 | PMF1 | protein_coding | 42.1574846 | 11.39668295 |
| ENSG00000172809.12 | RPL38 | protein_coding | 42.16586413 | 17.95083061 |
| ENSG00000078369.17 | GNB1 | protein_coding | 42.59092923 | 13.28597187 |
| ENSG00000182899.14 | RPL35A | protein_coding | 42.65700068 | 17.68384982 |
| ENSG00000198952.8 | SMG5 | protein_coding | 42.78118935 | 12.44366486 |
| ENSG00000175634.14 | RPS6KB2 | protein_coding | 43.12536946 | 8.113000618 |
| ENSG00000243746.1 | EEF1A1P10 | processed_pseudogene | 43.18782367 | 1.487616667 |
| ENSG00000117984.12 | CTSD | protein_coding | 43.30089637 | 20.25741091 |
| ENSG00000197756.9 | RPL37A | protein_coding | 43.4360163 | 11.44718514 |
| ENSG00000104964.14 | AES | protein_coding | 43.69707005 | 19.66510914 |
| ENSG00000172270.18 | BSG | protein_coding | 43.98838637 | 16.86102407 |
| ENSG00000172270.18 | BSG | protein_coding | 44.01884423 | 19.00298438 |
| ENSG00000269821.1 | KCNQ1OT1 | antisense | 44.20323945 | 6.512679448 |
| ENSG00000201390.1 | RNU6-1141P | snRNA | 44.34592775 | 7.231891557 |
| ENSG00000103495.13 | MAZ | protein_coding | 44.60061646 | 21.350976 |
| ENSG00000196821.9 | C6orf106 | protein_coding | 45.20298599 | 12.61073367 |
| ENSG00000185624.14 | P4HB | protein_coding | 45.30450762 | 20.37664369 |
| ENSG00000079313.11 | REXO1 | protein_coding | 45.42294948 | 10.77727925 |
| ENSG00000120885.19 | CLU | protein_coding | 45.63673028 | 16.49690103 |
| ENSG00000185883.10 | ATP6V0C | protein_coding | 45.83846506 | 16.88828646 |
| ENSG00000260272.1 | RP11-20I23.1 | protein_coding | 45.83846506 | 16.88828646 |
| ENSG00000162736.15 | NCSTN | protein_coding | 46.17819626 | 12.97877572 |
| ENSG00000114867.19 | EIF4G1 | protein_coding | 46.22921131 | 19.05734868 |
| ENSG00000245848.2 | CEBPA | protein_coding | 46.46535194 | 15.51990257 |
| ENSG00000084234.16 | APLP2 | protein_coding | 46.81849129 | 15.29966542 |
| ENSG00000015676.17 | NUDCD3 | protein_coding | 47.26932237 | 9.100446365 |
| ENSG00000198952.8 | SMG5 | protein_coding | 47.48342097 | 14.99774469 |
| ENSG00000116133.11 | DHCR24 | protein_coding | 47.80840758 | 16.38419653 |
| ENSG00000141522.11 | ARHGDIA | protein_coding | 47.89252982 | 11.00529753 |
| ENSG00000167508.10 | MVD | protein_coding | 48.18057623 | 12.52393513 |
| ENSG00000163220.10 | S100A9 | protein_coding | 48.41641641 | 23.22219696 |
| ENSG00000168137.15 | SETD5 | protein_coding | 48.43972523 | 16.81052189 |
| ENSG00000123144.10 | C19orf43 | protein_coding | 48.79176198 | 13.48939392 |
| ENSG00000101421.3 | CHMP4B | protein_coding | 48.84410288 | 14.59591199 |
| ENSG00000104969.9 | SGTA | protein_coding | 48.88487455 | 15.74139257 |
| ENSG00000111057.10 | KRT18 | protein_coding | 49.41381015 | 15.43169072 |
| ENSG00000101160.13 | CTSZ | protein_coding | 49.63323181 | 13.33511763 |
| ENSG00000132507.17 | EIF5A | protein_coding | 49.94115057 | 24.04627556 |

| | | | | |
|---|---|---|---|---|
| ENSG00000172270.18 | BSG | protein_coding | 50.45461664 | 23.31542021 |
| ENSG00000105373.18 | GLTSCR2 | protein_coding | 50.54102743 | 14.16196596 |
| ENSG00000197746.13 | PSAP | protein_coding | 50.99624759 | 16.75349143 |
| ENSG00000105722.9 | ERF | protein_coding | 51.13406765 | 12.12351573 |
| ENSG00000117984.12 | CTSD | protein_coding | 51.14733436 | 23.7638361 |
| ENSG00000171858.17 | RPS21 | protein_coding | 51.44933977 | 13.8301167 |
| ENSG00000142798.16 | HSPG2 | protein_coding | 51.45368511 | 10.86533065 |
| ENSG00000242590.1 | RP11-54O7.14 | sense_intronic | 51.46778192 | 22.54570592 |
| ENSG00000128272.14 | ATF4 | protein_coding | 51.8758488 | 18.71151093 |
| ENSG00000139645.9 | ANKRD52 | protein_coding | 51.97289718 | 13.08772167 |
| ENSG00000089597.16 | GANAB | protein_coding | 52.03262174 | 21.86551321 |
| ENSG00000111737.11 | RAB35 | protein_coding | 52.26599339 | 10.03221807 |
| ENSG00000173113.6 | TRMT112 | protein_coding | 52.59568169 | 19.1545945 |
| ENSG00000173120.14 | KDM2A | protein_coding | 52.64430921 | 24.61517708 |
| ENSG00000159335.15 | PTMS | protein_coding | 52.70408144 | 25.14000111 |
| ENSG00000130203.9 | APOE | protein_coding | 53.0212992 | 15.88402561 |
| ENSG00000101439.8 | CST3 | protein_coding | 53.0234272 | 22.49395088 |
| ENSG00000173120.14 | KDM2A | protein_coding | 53.19996735 | 26.01332905 |
| ENSG00000200674.1 | RN7SKP160 | misc_RNA | 53.21720999 | 12.01473023 |
| ENSG00000101444.12 | AHCY | protein_coding | 53.25439915 | 23.44243411 |
| ENSG00000084234.16 | APLP2 | protein_coding | 53.35589204 | 20.48668204 |
| ENSG00000109475.16 | RPL34 | protein_coding | 54.16670273 | 24.37420582 |
| ENSG00000105701.15 | FKBP8 | protein_coding | 54.64627604 | 8.547107102 |
| ENSG00000130175.9 | PRKCSH | protein_coding | 54.8160743 | 20.11869688 |
| ENSG00000173020.10 | ADRBK1 | protein_coding | 55.44676233 | 20.2352635 |
| ENSG00000264364.2 | DYNLL2 | protein_coding | 55.59573847 | 16.07574753 |
| ENSG00000135390.17 | ATP5G2 | protein_coding | 56.00226686 | 23.53315166 |
| ENSG00000135390.17 | ATP5G2 | protein_coding | 56.23637405 | 23.51367041 |
| ENSG00000091542.8 | ALKBH5 | protein_coding | 56.39542339 | 22.33842175 |
| ENSG00000115053.15 | NCL | protein_coding | 56.8424388 | 16.66538315 |
| ENSG00000280087.1 | CTB-129P6.7 | TEC | 56.85035449 | 24.30422238 |
| ENSG00000068028.17 | RASSF1 | protein_coding | 56.98577565 | 15.94230892 |
| ENSG00000113966.9 | ARL6 | protein_coding | 57.18969668 | 1.207682898 |
| ENSG00000123144.10 | C19orf43 | protein_coding | 57.38838479 | 18.89931385 |
| ENSG00000120885.19 | CLU | protein_coding | 58.47750022 | 28.72283444 |
| ENSG00000200246.1 | RNA5SP263 | rRNA | 58.91610421 | 1.417633225 |
| ENSG00000167792.11 | NDUFV1 | protein_coding | 59.28495242 | 25.48464289 |
| ENSG00000234906.8 | APOC2 | protein_coding | 59.31880428 | 25.58188872 |
| ENSG00000172757.12 | CFL1 | protein_coding | 59.49707664 | 28.80963298 |
| ENSG00000176340.3 | COX8A | protein_coding | 59.97468694 | 25.45487481 |
| ENSG00000184640.17 | Sep.09 | protein_coding | 60.03753615 | 23.71317345 |
| ENSG00000126012.11 | KDM5C | protein_coding | 60.13922197 | 25.75412938 |
| ENSG00000115053.15 | NCL | protein_coding | 60.21130054 | 24.7381684 |
| ENSG00000134824.13 | FADS2 | protein_coding | 60.72107139 | 28.31724319 |
| ENSG00000089693.10 | MLF2 | protein_coding | 60.78816824 | 22.81927187 |
| ENSG00000119383.19 | PPP2R4 | protein_coding | 60.80395793 | 16.89606758 |
| ENSG00000252431.1 | RNU6-1247P | snRNA | 61.77926578 | 1.27766634 |
| ENSG00000174903.14 | RAB1B | protein_coding | 62.16988414 | 25.69709892 |
| ENSG00000197746.13 | PSAP | protein_coding | 62.34514014 | 30.29338753 |
| ENSG00000252744.1 | Y_RNA | misc_RNA | 63.66210964 | 5.250632245 |
| ENSG00000149480.6 | MTA2 | protein_coding | 63.74196687 | 13.33637048 |
| ENSG00000197746.13 | PSAP | protein_coding | 63.86169906 | 29.99924793 |

| | | | | |
|---|---|---|---|---|
| ENSG00000165527.6 | ARF6 | protein_coding | 64.39434961 | 22.50298486 |
| ENSG00000115268.9 | RPS15 | protein_coding | 64.46967916 | 22.25673817 |
| ENSG00000118137.9 | APOA1 | protein_coding | 64.56411599 | 25.98356097 |
| ENSG00000102119.10 | EMD | protein_coding | 65.08961905 | 15.02876562 |
| ENSG00000106927.11 | AMBP | protein_coding | 65.16838428 | 31.80951943 |
| ENSG00000092841.18 | MYL6 | protein_coding | 65.24302517 | 31.31321063 |
| ENSG00000143321.18 | HDGF | protein_coding | 65.26253961 | 22.91761008 |
| ENSG00000280118.1 | RP11-385H1.1 | TEC | 65.27794489 | 0 |
| ENSG00000201822.1 | RNA5SP149 | rRNA | 65.41849146 | 17.62290036 |
| ENSG00000130203.9 | APOE | protein_coding | 65.74773945 | 23.69777166 |
| ENSG00000139645.9 | ANKRD52 | protein_coding | 65.86462244 | 10.22127383 |
| ENSG00000115268.9 | RPS15 | protein_coding | 65.89653923 | 32.20476697 |
| ENSG00000155506.16 | LARP1 | protein_coding | 67.20952681 | 20.77831594 |
| ENSG00000173599.13 | PC | protein_coding | 67.32570366 | 24.8251274 |
| ENSG00000047849.21 | MAP4 | protein_coding | 67.49687543 | 18.01428578 |
| ENSG00000198858.9 | R3HDM4 | protein_coding | 67.53917343 | 25.13347283 |
| ENSG00000130829.17 | DUSP9 | protein_coding | 68.45427091 | 27.5346328 |
| ENSG00000117984.12 | CTSD | protein_coding | 68.62732251 | 26.57836844 |
| 26992 | 26992 | tRNAscan | 69.34050728 | 22.33064062 |
| ENSG00000002834.17 | LASP1 | protein_coding | 69.7100781 | 21.80717301 |
| ENSG00000185624.14 | P4HB | protein_coding | 69.71186935 | 30.65233872 |
| ENSG00000213593.9 | TMX2 | protein_coding | 72.38466391 | 32.65819425 |
| ENSG00000201428.1 | RN7SKP71 | misc_RNA | 72.99284988 | 1.789537392 |
| ENSG00000160789.19 | LMNA | protein_coding | 73.24217541 | 30.32190276 |
| ENSG00000126247.10 | CAPNS1 | protein_coding | 73.30699901 | 34.39973516 |
| ENSG00000110492.15 | MDK | protein_coding | 73.60838618 | 27.98163538 |
| ENSG00000188846.13 | RPL14 | protein_coding | 73.93989683 | 32.61161107 |
| ENSG00000111678.10 | C12orf57 | protein_coding | 74.34672027 | 24.92618866 |
| ENSG00000258674.5 | CTC-260F20.3 | protein_coding | 74.44178828 | 33.72340457 |
| ENSG00000132507.17 | EIF5A | protein_coding | 74.48671 | 29.00788318 |
| ENSG00000186010.18 | NDUFA13 | protein_coding | 74.63296786 | 31.81338155 |
| ENSG00000197746.13 | PSAP | protein_coding | 74.76092295 | 27.5553669 |
| ENSG00000071553.16 | ATP6AP1 | protein_coding | 75.07550719 | 23.62000709 |
| ENSG00000130175.9 | PRKCSH | protein_coding | 75.21876605 | 29.75435767 |
| ENSG00000125991.18 | ERGIC3 | protein_coding | 75.66837487 | 31.05912593 |
| ENSG00000131871.14 | VIMP | protein_coding | 75.6821122 | 5.73006906 |
| ENSG00000163348.3 | PYGO2 | protein_coding | 75.7749668 | 21.6787458 |
| ENSG00000173113.6 | TRMT112 | protein_coding | 75.91311525 | 27.07451678 |
| ENSG00000109072.13 | VTN | protein_coding | 76.03494512 | 29.59365668 |
| ENSG00000238923.1 | RNU7-1 | snRNA | 76.29916962 | 25.97191772 |
| ENSG00000164190.16 | NIPBL | protein_coding | 76.663714 | 1.045729058 |
| ENSG00000118137.9 | APOA1 | protein_coding | 77.52222519 | 34.85985117 |
| ENSG00000237729.2 | AC002075.4 | processed_pseudogene | 77.69211124 | 1.1157125 |
| ENSG00000120885.19 | CLU | protein_coding | 78.09766269 | 27.59558227 |
| ENSG00000172757.12 | CFL1 | protein_coding | 78.39593276 | 33.35008709 |
| ENSG00000173762.7 | CD7 | protein_coding | 79.24178805 | 29.00929649 |
| ENSG00000089597.16 | GANAB | protein_coding | 79.55495189 | 22.20373029 |
| ENSG00000160867.14 | FGFR4 | protein_coding | 79.69355438 | 30.42943541 |
| ENSG00000211450.9 | C11orf31 | protein_coding | 79.87479779 | 27.12386969 |
| ENSG00000178971.13 | CTC1 | protein_coding | 80.35989769 | 20.25088264 |
| ENSG00000105722.9 | ERF | protein_coding | 81.28407849 | 26.00946693 |
| ENSG00000167986.13 | DDB1 | protein_coding | 82.42332962 | 31.2678803 |

| | | | | |
|---|---|---|---|---|
| ENSG00000072518.20 | MARK2 | protein_coding | 82.99516724 | 26.90227612 |
| ENSG00000184640.17 | Sep.09 | protein_coding | 84.33755305 | 37.80113339 |
| ENSG00000130733.10 | YIPF2 | protein_coding | 84.505573 | 27.07718294 |
| ENSG00000173020.10 | ADRBK1 | protein_coding | 85.79068004 | 32.20993883 |
| ENSG00000089597.16 | GANAB | protein_coding | 85.88101764 | 35.32122004 |
| ENSG00000175061.17 | LRRC75A-AS1 | processed_transcript | 85.90921429 | 25.16590707 |
| ENSG00000159335.15 | PTMS | protein_coding | 86.37026601 | 42.54777928 |
| ENSG00000125691.12 | RPL23 | protein_coding | 87.84585356 | 38.20312656 |
| ENSG00000100836.10 | PABPN1 | protein_coding | 87.9341282 | 28.38597378 |
| ENSG00000110492.15 | MDK | protein_coding | 88.05682291 | 36.05975295 |
| ENSG00000104964.14 | AES | protein_coding | 88.88114926 | 33.05083253 |
| ENSG00000166681.13 | NGFRAP1 | protein_coding | 89.54003248 | 34.66160098 |
| ENSG00000133065.10 | SLC41A1 | protein_coding | 90.41351288 | 2.113445071 |
| ENSG00000108107.12 | RPL28 | protein_coding | 90.96644911 | 43.73755469 |
| ENSG00000130165.10 | ELOF1 | protein_coding | 91.00655184 | 14.47292066 |
| ENSG00000089693.10 | MLF2 | protein_coding | 91.28588942 | 36.3758795 |
| ENSG00000188157.13 | AGRN | protein_coding | 91.30469106 | 37.9463757 |
| ENSG00000188186.10 | LAMTOR4 | protein_coding | 91.91345437 | 28.28622226 |
| ENSG00000146830.9 | GIGYF1 | protein_coding | 91.97054099 | 35.85372163 |
| ENSG00000102858.12 | MGRN1 | protein_coding | 94.20002672 | 25.32143621 |
| ENSG00000142534.6 | RPS11 | protein_coding | 94.22030309 | 46.29272691 |
| ENSG00000089248.6 | ERP29 | protein_coding | 95.09807753 | 30.83502666 |
| ENSG00000118137.9 | APOA1 | protein_coding | 96.82734557 | 36.3901889 |
| ENSG00000167526.13 | RPL13 | protein_coding | 98.21754252 | 45.91178877 |
| ENSG00000101439.8 | CST3 | protein_coding | 99.65957248 | 35.98073553 |
| ENSG00000186010.18 | NDUFA13 | protein_coding | 100.4661348 | 41.46977645 |
| ENSG00000092841.18 | MYL6 | protein_coding | 100.6052208 | 47.04431637 |
| ENSG00000167508.10 | MVD | protein_coding | 100.8474745 | 26.42795427 |
| ENSG00000167470.12 | MIDN | protein_coding | 100.9429914 | 47.26847253 |
| ENSG00000224916.8 | APOC4-APOC2 | protein_coding | 101.8622668 | 44.33747713 |
| ENSG00000047849.21 | MAP4 | protein_coding | 102.526831 | 36.5909448 |
| ENSG00000184840.11 | TMED9 | protein_coding | 104.0367608 | 47.1544116 |
| ENSG00000137309.19 | HMGA1 | protein_coding | 104.1639756 | 32.96262068 |
| ENSG00000168159.10 | RNF187 | protein_coding | 106.0120944 | 33.58877 |
| ENSG00000047849.21 | MAP4 | protein_coding | 106.408452 | 38.23665318 |
| ENSG00000161011.19 | SQSTM1 | protein_coding | 106.7048598 | 41.19115241 |
| ENSG00000198952.8 | SMG5 | protein_coding | 106.941518 | 36.01050361 |
| ENSG00000092841.18 | MYL6 | protein_coding | 107.1104307 | 49.55039971 |
| ENSG00000115268.9 | RPS15 | protein_coding | 109.4597849 | 52.66810563 |
| ENSG00000132507.17 | EIF5A | protein_coding | 110.2215798 | 42.38196332 |
| ENSG00000114942.13 | EEF1B2 | protein_coding | 110.4481445 | 49.21996376 |
| ENSG00000130203.9 | APOE | protein_coding | 110.7728602 | 44.59923938 |
| ENSG00000186010.18 | NDUFA13 | protein_coding | 110.9418025 | 49.13175191 |
| ENSG00000151176.7 | PLBD2 | protein_coding | 112.4554723 | 41.37274797 |
| ENSG00000070756.13 | PABPC1 | protein_coding | 112.6039739 | 42.91331563 |
| ENSG00000092841.18 | MYL6 | protein_coding | 115.2391017 | 36.94211486 |
| ENSG00000138107.11 | ACTR1A | protein_coding | 118.1001851 | 44.28681449 |
| ENSG00000172757.12 | CFL1 | protein_coding | 119.0673147 | 52.01762412 |
| ENSG00000245848.2 | CEBPA | protein_coding | 119.7944399 | 43.73989993 |
| ENSG00000117984.12 | CTSD | protein_coding | 120.9900988 | 47.96569768 |
| ENSG00000105722.9 | ERF | protein_coding | 121.4846058 | 22.93056306 |
| ENSG00000110721.11 | CHKA | protein_coding | 124.3449256 | 53.47270298 |

| | | | | |
|---|---|---|---|---|
| ENSG00000103495.13 | MAZ | protein_coding | 125.2044584 | 44.93865243 |
| ENSG00000198911.11 | SREBF2 | protein_coding | 125.3658144 | 34.99720879 |
| ENSG00000159335.15 | PTMS | protein_coding | 125.6192194 | 50.47558626 |
| ENSG00000253626.3 | EIF5AL1 | protein_coding | 126.9180308 | 33.65326067 |
| ENSG00000210049.1 | MT-TF | Mt_tRNA | 127.1563705 | 53.10999324 |
| ENSG00000051523.10 | CYBA | protein_coding | 127.9704465 | 56.15771572 |
| ENSG00000145592.13 | RPL37 | protein_coding | 128.0380528 | 56.12402863 |
| ENSG00000159335.15 | PTMS | protein_coding | 129.7893212 | 31.4868646 |
| ENSG00000130255.12 | RPL36 | protein_coding | 131.241099 | 62.53319801 |
| ENSG00000184897.5 | H1FX | protein_coding | 131.2742578 | 32.90710709 |
| ENSG00000115268.9 | RPS15 | protein_coding | 132.1739113 | 48.51386509 |
| ENSG00000154277.12 | UCHL1 | protein_coding | 132.7879378 | 46.9107173 |
| ENSG00000155366.16 | RHOC | protein_coding | 138.2031561 | 57.04149094 |
| ENSG00000160867.14 | FGFR4 | protein_coding | 138.4253709 | 59.82353216 |
| ENSG00000034510.5 | TMSB10 | protein_coding | 140.5776405 | 69.34252275 |
| ENSG00000196465.10 | MYL6B | protein_coding | 140.8880617 | 66.44254829 |
| ENSG00000159335.15 | PTMS | protein_coding | 140.9700433 | 36.60013923 |
| ENSG00000168066.20 | SF1 | protein_coding | 141.050985 | 55.55757593 |
| ENSG00000118137.9 | APOA1 | protein_coding | 141.1278038 | 44.64832826 |
| ENSG00000088247.15 | KHSRP | protein_coding | 141.3477886 | 62.62239868 |
| ENSG00000198561.12 | CTNND1 | protein_coding | 141.5447134 | 66.71845948 |
| ENSG00000172270.18 | BSG | protein_coding | 144.3044326 | 66.91410041 |
| ENSG00000174903.14 | RAB1B | protein_coding | 145.0876655 | 62.64198351 |
| ENSG00000130725.7 | UBE2M | protein_coding | 145.2765848 | 63.85483826 |
| ENSG00000168066.20 | SF1 | protein_coding | 151.0131571 | 49.94031494 |
| ENSG00000104852.14 | SNRNP70 | protein_coding | 157.5973457 | 57.19685961 |
| ENSG00000167792.11 | NDUFV1 | protein_coding | 158.6539623 | 61.92397756 |
| ENSG00000178252.17 | WDR6 | protein_coding | 162.8547459 | 45.77307474 |
| ENSG00000089597.16 | GANAB | protein_coding | 164.3823338 | 59.79773997 |
| ENSG00000197746.13 | PSAP | protein_coding | 167.5955932 | 74.06196315 |
| ENSG00000110492.15 | MDK | protein_coding | 167.9196625 | 75.08548791 |
| ENSG00000159335.15 | PTMS | protein_coding | 173.425188 | 54.17122766 |
| ENSG00000167685.14 | ZNF444 | protein_coding | 174.5858606 | 42.40922571 |
| ENSG00000213741.8 | RPS29 | protein_coding | 177.7231404 | 75.87233822 |
| ENSG00000085733.15 | CTTN | protein_coding | 178.1275197 | 54.14636741 |
| ENSG00000159335.15 | PTMS | protein_coding | 178.650862 | 69.74043645 |
| ENSG00000167815.11 | PRDX2 | protein_coding | 179.5197203 | 51.74928691 |
| ENSG00000130203.9 | APOE | protein_coding | 179.6126467 | 70.43379949 |
| ENSG00000130402.11 | ACTN4 | protein_coding | 184.3825806 | 64.73720017 |
| ENSG00000169241.17 | SLC50A1 | protein_coding | 184.9955183 | 72.87893336 |
| ENSG00000184840.11 | TMED9 | protein_coding | 186.1254227 | 85.48574803 |
| ENSG00000204628.11 | GNB2L1 | protein_coding | 186.174549 | 59.93117858 |
| ENSG00000116251.9 | RPL22 | protein_coding | 188.5400014 | 70.45578644 |
| **ENSG00000008988.9** | **RPS20** | **protein_coding** | **188.9078877** | **74.64125506** |
| ENSG00000109062.9 | SLC9A3R1 | protein_coding | 193.1839235 | 78.01914946 |
| ENSG00000172270.18 | BSG | protein_coding | 194.8935032 | 87.6950825 |
| ENSG00000125691.12 | RPL23 | protein_coding | 208.5181321 | 97.85942945 |
| ENSG00000167978.16 | SRRM2 | protein_coding | 211.1766838 | 96.66856165 |
| ENSG00000140264.19 | SERF2 | protein_coding | 212.1451219 | 88.57901818 |
| ENSG00000210164.1 | MT-TG | Mt_tRNA | 212.8443778 | 78.34337806 |
| ENSG00000110700.6 | RPS13 | protein_coding | 216.1771652 | 78.76588798 |
| ENSG00000130203.9 | APOE | protein_coding | 216.2113364 | 89.29692055 |

| | | | | |
|---|---|---|---|---|
| ENSG00000110492.15 | MDK | protein_coding | 217.925044 | 93.5949901 |
| ENSG00000177954.11 | RPS27 | protein_coding | 225.0556517 | 87.81447574 |
| ENSG00000167685.14 | ZNF444 | protein_coding | 227.0221078 | 86.36152477 |
| ENSG00000120885.19 | CLU | protein_coding | 231.3876277 | 96.77119667 |
| ENSG00000034510.5 | TMSB10 | protein_coding | 231.9089162 | 101.7375983 |
| ENSG00000266173.5 | STRADA | protein_coding | 233.7134291 | 4.676558877 |
| ENSG00000221983.7 | UBA52 | protein_coding | 236.6417797 | 79.95131989 |
| ENSG00000161016.15 | RPL8 | protein_coding | 242.1957247 | 57.07261544 |
| ENSG00000063177.12 | RPL18 | protein_coding | 243.6485399 | 106.8459185 |
| ENSG00000149273.14 | RPS3 | protein_coding | 243.7766017 | 89.5507879 |
| ENSG00000103495.13 | MAZ | protein_coding | 251.1478867 | 99.97511618 |
| ENSG00000231221.1 | LINC01593 | lincRNA | 267.752343 | 5.040681918 |
| ENSG00000068400.13 | GRIPAP1 | protein_coding | 271.750653 | 71.46130014 |
| ENSG00000197746.13 | PSAP | protein_coding | 273.0996006 | 117.3732494 |
| ENSG00000130829.17 | DUSP9 | protein_coding | 275.4352725 | 116.8645986 |
| ENSG00000136942.14 | RPL35 | protein_coding | 288.8904239 | 106.9935629 |
| ENSG00000034510.5 | TMSB10 | protein_coding | 290.135015 | 121.4766566 |
| ENSG00000105193.8 | RPS16 | protein_coding | 291.9042885 | 114.8735338 |
| ENSG00000184840.11 | TMED9 | protein_coding | 292.0115341 | 123.6279251 |
| ENSG00000172809.12 | RPL38 | protein_coding | 292.8224508 | 134.4662005 |
| ENSG00000063177.12 | RPL18 | protein_coding | 294.562508 | 90.73769001 |
| ENSG00000168028.13 | RPSA | protein_coding | 296.5645135 | 99.27451029 |
| ENSG00000167685.14 | ZNF444 | protein_coding | 298.3857265 | 109.4609478 |
| ENSG00000118137.9 | APOA1 | protein_coding | 319.2196354 | 119.5978719 |
| ENSG00000070756.13 | PABPC1 | protein_coding | 331.4286445 | 153.0298599 |
| ENSG00000083845.8 | RPS5 | protein_coding | 337.4141102 | 92.66980355 |
| ENSG00000124614.13 | RPS10 | protein_coding | 343.0944248 | 118.0833137 |
| **ENSG00000137154.12** | **RPS6** | **protein_coding** | **353.6709997** | **86.99540854** |
| ENSG00000186468.12 | RPS23 | protein_coding | 407.9755422 | 177.8280458 |
| ENSG00000210100.1 | MT-TI | Mt_tRNA | 420.7436523 | 194.4813612 |
| ENSG00000134824.13 | FADS2 | protein_coding | 421.4092296 | 153.5968667 |
| ENSG00000138326.18 | RPS24 | protein_coding | 427.38332 | 150.4682163 |
| ENSG00000200434.1 | RNA5-8SP2 | rRNA | 435.5737463 | 31.00471494 |
| ENSG00000173020.10 | ADRBK1 | protein_coding | 440.7611132 | 167.2804882 |
| ENSG00000108107.12 | RPL28 | protein_coding | 443.3665415 | 194.3063508 |
| ENSG00000089248.6 | ERP29 | protein_coding | 453.4432046 | 150.5159954 |
| ENSG00000132507.17 | EIF5A | protein_coding | 455.2918635 | 175.026917 |
| ENSG00000179218.13 | CALR | protein_coding | 456.4946832 | 136.3092374 |
| ENSG00000167978.16 | SRRM2 | protein_coding | 460.2328079 | 135.2069751 |
| ENSG00000125844.15 | RRBP1 | protein_coding | 462.945466 | 163.2340546 |
| ENSG00000142534.6 | RPS11 | protein_coding | 480.5504944 | 178.1814007 |
| ENSG00000130203.9 | APOE | protein_coding | 487.4170256 | 180.4273991 |
| ENSG00000197746.13 | PSAP | protein_coding | 509.5433388 | 196.303276 |
| ENSG00000110107.8 | PRPF19 | protein_coding | 515.9945291 | 203.9270035 |
| **ENSG00000147257.13** | **GPC3** | **protein_coding** | **532.864386** | **178.1531028** |
| ENSG00000089597.16 | GANAB | protein_coding | 535.0646472 | 231.706076 |
| ENSG00000167244.17 | IGF2 | protein_coding | 559.5054613 | 195.36378 |
| ENSG00000159335.15 | PTMS | protein_coding | 610.1681404 | 254.4473558 |
| ENSG00000241553.12 | ARPC4 | protein_coding | 610.5465403 | 282.5139621 |
| ENSG00000210144.1 | MT-TY | Mt_tRNA | 614.9614602 | 272.3644059 |
| ENSG00000197756.9 | RPL37A | protein_coding | 630.08485 | 249.312736 |
| ENSG00000105372.6 | RPS19 | protein_coding | 634.749785 | 233.2005419 |

| | | | | |
|---|---|---|---|---|
| **ENSG00000167526.13** | **RPL13** | **protein_coding** | **641.3875677** | **216.2224698** |
| ENSG00000105372.6 | RPS19 | protein_coding | 685.8776829 | 306.7659789 |
| ENSG00000167244.17 | IGF2 | protein_coding | 690.1388499 | 254.4988103 |
| ENSG00000130203.9 | APOE | protein_coding | 719.0813302 | 239.8549224 |
| ENSG00000089597.16 | GANAB | protein_coding | 719.8451419 | 270.0346996 |
| **ENSG00000117984.12** | **CTSD** | **protein_coding** | **843.3342901** | **274.0704528** |
| ENSG00000112306.7 | RPS12 | protein_coding | 970.608266 | 408.9356127 |
| ENSG00000177600.8 | RPLP2 | protein_coding | 1003.366686 | 292.5724639 |
| ENSG00000070756.13 | PABPC1 | protein_coding | 2328.056117 | 720.0658393 |
